# Monomeric amyloid-β inhibits microglial inflammatory activity in the brain via an APP/heterotrimeric G protein-mediated pathway

**DOI:** 10.1101/2023.07.24.550398

**Authors:** Hyo Jun Kwon, Devi Santhosh, Zhen Huang

## Abstract

Microglia, the resident immune cell of the brain, play critical roles in brain development, function, and disease. However, how microglial activity is regulated in this process remains to be elucidated. Here we report an amyloid precursor protein (APP) and heterotrimeric G protein-mediated pathway that negatively regulates microglial inflammatory activation during cerebral cortex development. Disruption of this pathway results in dysregulated microglial activity, excessive extracellular matrix proteinase production, cortical basement membrane breach, and laminar assembly disruption. We further show that this pathway is activated by amyloid β (Aβ), the cleavage product of APP that accumulates in large quantities as plaques in the Alzheimer’s disease brain. Specifically, we find Aβ monomers potently suppress inflammatory cytokine transcription and secretion by brain microglia, in an APP and heterotrimeric G protein-dependent manner. These results discover a previously unknown activity of Aβ as a negative regulator of brain microglia as well as a new pathway that mediates the signal transduction. They shed new light on the cell-cell communication mechanisms that regulate brain immune homeostasis and may facilitate further insight into Alzheimer’s disease pathogenesis.

## INTRODUCTION

The cerebral cortex is well known for its exquisite six-layered architecture assembled from an extensive cohort of neural cell types during development. These cells form the foundation of the brain neural circuitry and include locally born excitatory neurons that populate the cortex through radial migration as well as inhibitory neurons born outside that integrate into the cortex through tangential migration. These cells also include a large number of astroglial cells and oligodendrocytes that are born both inside and outside the cortex. Decades of studies have elucidated elaborate cell-cell signaling mechanisms that coordinate the interactions of these cells during development, mechanisms that ensure the precise assembly of the cortical laminar architecture and the brain neural circuitry (Franco and Muller, 2013; Hansen et al., 2011; Marin and Rubenstein, 2003). Besides neural lineage cells, however, the brain also contains many non-neural cells. Microglia, for example, are the main bona fide immune cell type of the brain parenchyma and constitute 10-15% of all brain cells. It remains poorly understood how microglial behavior and activity is regulated during their integration into the brain.

Microglia are unique among brain cell types in that they are of a myeloid lineage closely related to macrophages in the periphery. They originate from outside the nervous system and integrate into the cerebral cortex during early development at the onset of laminar assembly (Ginhoux et al., 2010; Hattori et al., 2023). Microglia play crucial roles in both brain health and disease throughout life (Colonna and Butovsky, 2017; Iwasawa et al., 2022; Li and Barres, 2018). As immune cells, they are well known to protect the brain against pathogens as well as damaged cells. They are also critically involved in many aspects of the development, function and plasticity of the brain neural circuitry. During aging, microglial activity has also been found to be among the most prominently affected and has been implicated in many neurodegenerative diseases (Shi and Holtzman, 2018; Yeh et al., 2017). As a myeloid lineage cell type, microglia are highly mobile and produce a large number of cytokines and chemokines that can in turn impact nearly all aspects of brain development, function, and plasticity. This raises the question of how their activity is regulated in brain immune homeostasis.

The amyloid precursor protein (APP) is a single-span transmembrane glycoprotein well known for mutations associated with the development of familial Alzheimer’s disease in humans (Selkoe and Hardy, 2016; Tanzi, 2012). APP can undergo series of proteolytic processing in its extracellular and transmembrane domains and produce a large number of cleavage products (De Strooper et al., 2012; Haass et al., 2012). Among them, amyloid β (Aβ), a product of about 40 amino acid residues is known to accumulate in large quantities in plaques in the brain of Alzheimer’s disease patients and forms one of the pathological hallmarks of Alzheimer’s disease. APP, however, can also transduce signals across the plasma membrane, both by activating cytoplasmic signaling partners and by transmitting cleavage products to the nucleus. Heterotrimeric G proteins, for example, have been found to interact with the APP cytoplasmic domain and implicated in APP regulated signaling in both invertebrates and mammals (Fogel et al., 2014; Milosch et al., 2014; Nishimoto et al., 1993; Ramaker et al., 2013). Heterotrimeric G protein function in turn depends critically on Ric8a, a guanine nucleotide exchange factor as well as a molecular chaperone essential for heterotrimeric G protein stability and activity (Gabay et al., 2011; Ma et al., 2012; Ma et al., 2017; Tall et al., 2003).

In this article, we report that APP and Ric8a-regulated heterotrimeric G proteins form a novel anti-inflammatory signaling pathway in microglia that prevents excessive microglial immune activation during cortical development. We show that this regulation is essential for the proper integration of microglia into the developing brain without disrupting the cortical architecture. In the absence of this pathway, microglia display an elevated reaction to immune stimulation, resulting in increased levels of extracellular matrix proteinases, breakdown of the cortical basement membrane, and disruption of proper neuronal layer formation. Furthermore, we show that this pathway is activated by monomeric Aβ, which strongly suppresses microglial inflammatory cytokine secretion and transcriptional activation, and that these effects depend on APP and Ric8a function in microglia. These results not only provide a potential new understanding of the normal biological function of Aβ but may also facilitate new insight into the pathogenesis of Alzheimer’s disease.

## RESULTS

### *emx1-cre*-mediated deletion of *ric8a* results in cortical neuron ectopia

In studies to determine the function of *ric8a* gene during corticogenesis, we deleted a conditional *ric8a* knockout allele using *emx1-cre,* a *cre* line designed to target dorsal forebrain neural progenitors and neurons beginning around embryonic day 10.5 (E10.5) (Gorski et al., 2002). We found this result in severe disruption of cortical organization (**Fig. 1A-G**). At both postnatal day 0 (P0) and 5 (P5), a large number of neuronal ectopia were observed at the pial surface. These ectopia are morphologically similar to those found in mouse models of cobblestone lissencephaly, including *dystroglycan* and related mutants, except that, in those mutants, the most severe ectopia were typically observed at the midline (Beggs et al., 2003; Belvindrah et al., 2006; Graus-Porta et al., 2001; Huang et al., 2006; Moore et al., 2002; Niewmierzycka et al., 2005; Satz et al., 2010). However, no ectopia were observed at the cortical midline in *ric8a/emx1-cre* mutants, which were exclusively located in the lateral cortex. These results indicate that *ric8a* plays a critical role in normal cortical development, but potentially by a mechanism distinct from the dystroglycan and related mutants.

**Figure 1.**
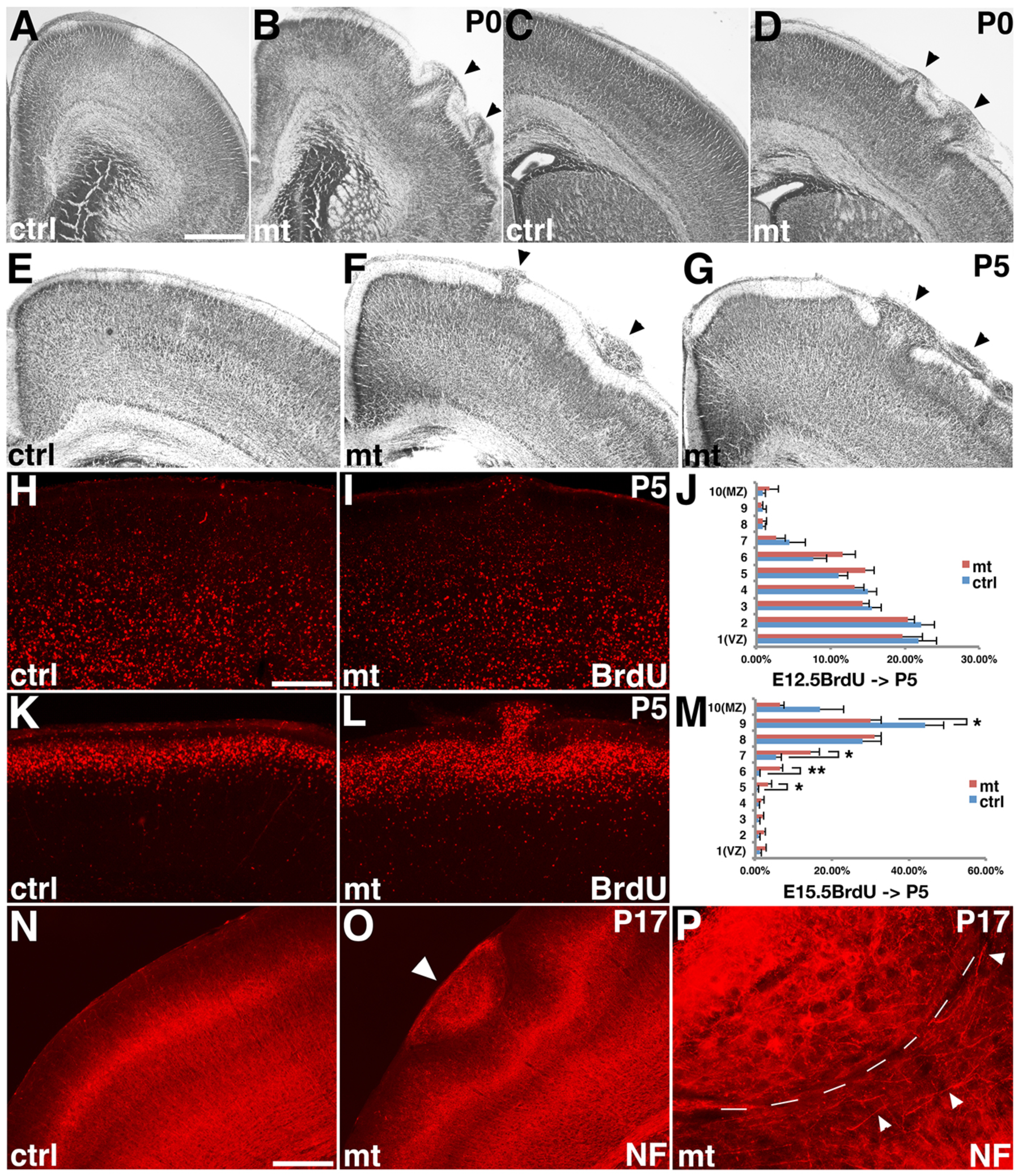
Deletion of *ric8a* using *emx1-cre* results in neuronal ectopia at the cortical pia. (**A** & **B**) Nissl staining of control (ctrl, **A**) and mutant (mt, **B**) anterior motor cortex at P0. See quantification of P0 ectopia in Fig. 3J & **K**. (**C** & **D**) Nissl staining of control (**C**) and mutant (**D**) somatosensory cortex at P0. See quantification of P0 ectopia in Fig. 7K & **L**. **(E-G**) Nissl staining of control (**E**) and different mutant (**F** & **G**) cortices at P5. (**H-J**) BrdU (in red) staining in control (**H**) and mutant (**I**) cortices at P5 after administration at E12.5. Quantification is shown in (**J**). No statistically significant differences were observed between control and mutant neurons in regions without ectopia. (**K-M**) BrdU staining in control (**K**) and mutant (**L**) cortices at P5 after administration at E15.5. Quantification is shown in (**M**). Neuronal migration appears slightly delayed in mutants as compared to controls. *, *P* < 0.05; **, *P* < 0.01; n = 5. (**N-P**) Neurofilament staining (NF, in red) of control (**N**) and mutant (**O**) cortices at P17. Dense ectopic nerve fibers were observed at ectopias in the mutant cortex (arrowhead in (**O**)), where neuronal processes (arrowheads in (**P**)) ran parallel to the border of ectopias (dash line in (**P**)). Scale bars, 640μm for (**A** & **B**), 400μm for (**C** & **D**), 500μm for (E-G), 375μm for (**H, I, K, L, N, O**), and 94μm for (**P**).

To determine how *ric8a* mutation results in cortical ectopia, we next determined the cellular composition of the ectopia. Through 5-bromo-2-deoxyuridine (BrdU)-based birthdating, we found that the ectopia consist of both early and late-born neurons (**Fig. 1H-M**), indicating ectopia formation embark during an early stage. Indeed, the ectopia stained positive for both Ctip2 and Cux1, transcriptional factors specific to lower and upper-layer cortical plate neurons, respectively (see Fig. S3). Interestingly, in cortical regions without ectopia, the radial migration of early-born neurons appeared to be largely normal (**Fig. 1J**), while the migration of late-born neurons appeared to be slightly delayed (**Fig. 1M**). The laminar targeting of Cux1 and Ctip2 positive neurons also appeared relatively normal in regions without ectopia (see Fig. S3), suggesting that the ectopia are unlikely to result from cell autonomous defects in cortical plate neurons. In support, we found that deletion of *ric8a* using a *nex-cre* (Goebbels et al., 2006), a *cre* line near-exclusively restricted to post-mitotic cortical neurons, did not result in any ectopia formation (data not shown). These results indicate that neuronal ectopia in *ric8a* mutants likely result from defects in cell types other than cortical plate neurons.

To determine the effects of *ric8a* mutation on cortical circuit assembly, we next examined the localization of other neuronal cell types as well as the development of synapses. We found that not only were excitatory neurons displaced in *ric8a* mutant cortices, but inhibitory neurons also migrated abnormally. For example, we found that in *ric8a* mutant cortices, parvalbumin-positive neurons failed to show their typical discrete layering, and many were far away from their normal locations (**Fig. S1A-B”**). The usual radial arrangement of the major processes of layer II/III neurons was also disorganized, most prominently in regions directly beneath the ectopia where the neuronal processes appeared to run in horizontal instead of the normal radial directions (**Fig. 1N-P**). At the synaptic level, the normal dense pattern of synaptic boutons in layer I was also perturbed, which showed instead a pattern typical of deeper layers in the mutants (**Fig. S1C-E’**). Thus, ectopia formation in *ric8a* mutants appears to severely disrupt the normal organization of neural circuits in the cerebral cortex.

### Basement membrane breach leads to ectopia formation during corticogenesis

To determine how the ectopia first develop, we next examined the *ric8a* mutant cortex at progressively earlier embryonic stages. At E16.5, we observed clear breaches in the pial basement membrane (**Fig. 2A-B”**), which indicates embryonic origin of the ectopia. Unlike classic models of cobblestone lissencephaly, where radial glial fibers typically retract from the pia, in *ric8a* mutants, the fibers instead extend beyond the breach sites. In regions without ectopia, we also observed a normal localization of Cajal-Retzius cells in the marginal zone (**Fig. 2C-E**). While missing at large ectopia (**Fig. 2D**), they still lined up relatively normally at small ectopia (**Fig. 2E**). We also observed normal expression of Reelin (**Fig. 2F&G**) as well as normal preplate splitting (**Fig. 2H&I**). This argues against a primary defect in the localization of Cajal-Retzius cells leading to ectopia formation. In line with this, we found that deletion of *ric8a* from Cajal-Retzius cells, using a *wnt3a-cre* (Yoshida et al., 2006), also did not result in ectopia (see Fig. 4). Thus, these results indicate that deficits in Cajal-Retzius cells are unlikely to be the primary cause leading to ectopia formation in *ric8a* mutants.

**Figure 2.**
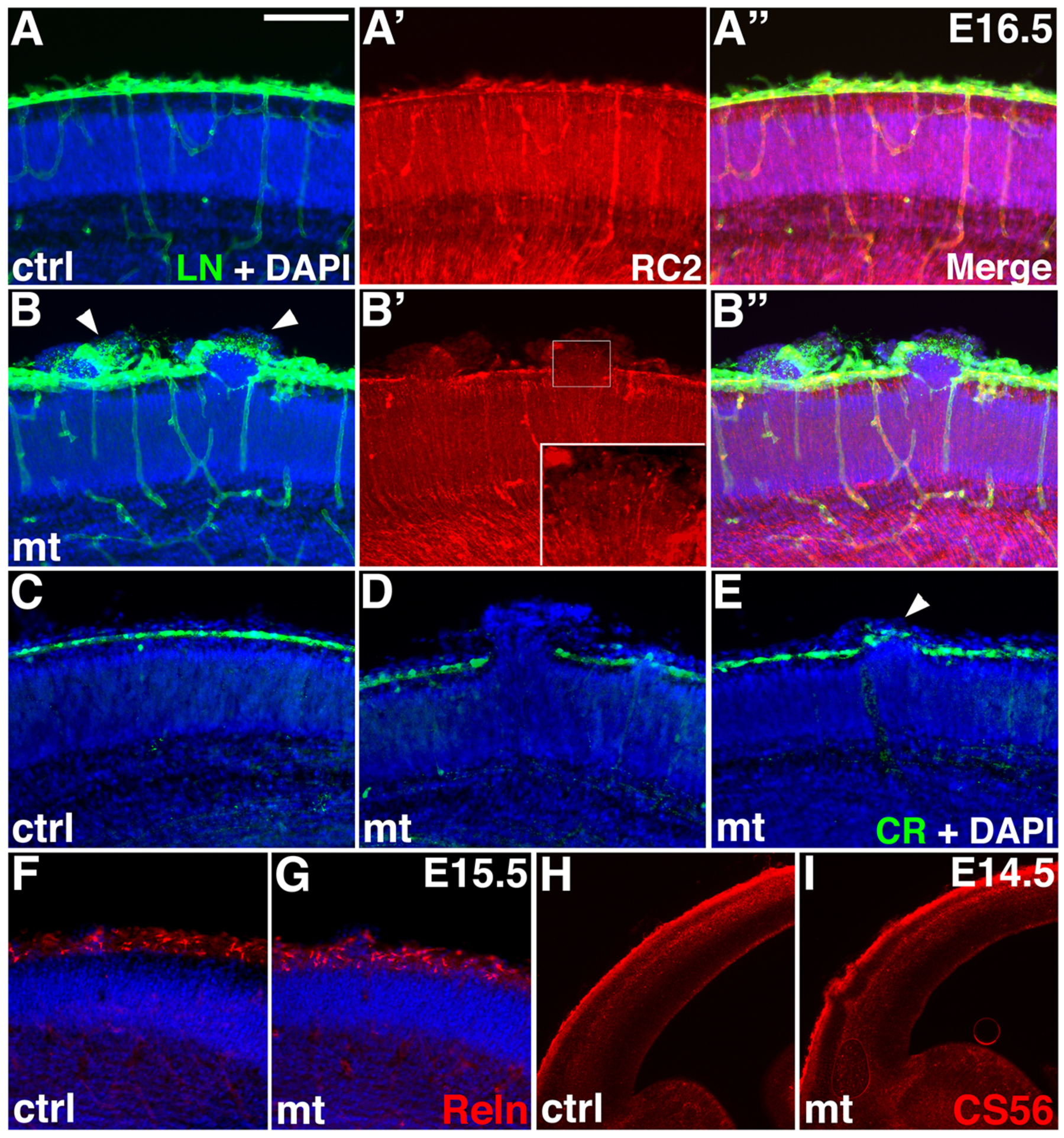
Neuronal ectopia in *ric8a* mutants result from pial basement membrane breach during embryogenesis. (**A-A”**) Laminin (LN, in green), radial glial marker RC2 (in red), and nuclear (DAPI, in blue) staining of control cortices at E16.5. A continuous basement membrane is observed at the pia, where radial glial endfeet are anchored. (**B**-**B”**) Laminin, RC2, and nuclear staining of *ric8a/emx1-cre* mutant cortices at E16.5. Neuronal ectopias are consistently observed at sites of basement membrane breakage (arrowheads in **B**). Radial glial fibers at these sites extend beyond the pia (inset in **B’**). (**C**-**E**) Calretinin (CR, in green) and nuclear (DAPI, in blue) staining of control (**C**) and mutant (**D** & **E**) cortices at E16.5. A continuous row of Calretinin positive Cajal-Retzius cells is observed in the marginal zone of control cortices (**C**). By contrast, in mutants, Cajal-Retzius cells are absent at large ectopias (**D**). However, they appear passively displaced by over-migrating neurons at small ectopias (arrowhead in **E**). **(F** & **G**) Reelin (Reln, in red) and nuclear (DAPI, in blue) staining of control (**F**) and *ric8a*/*emx1-cre* mutant (**G**) cortices at E15.5. Strong Reelin expression is observed in Cajal-Retzius cells in the marginal zone of both control and mutant cortices. **(H** & **I)** Chondroitin sulfate proteoglycan (CS56, in red) staining of control (**H**) and mutant (**I**) cortices at E14.5. Normal preplate splitting is observed in mutants. Scale bar in (**A**), 200μm for (**A**-**G)** and 500μm for (**H** & **I)**.

In classic cobblestone lissencephaly, neuronal ectopia result from primary defects in maintaining the integrity of pial basement membrane (Beggs et al., 2003; Graus-Porta et al., 2001; Moore et al., 2002; Satz et al., 2010). To determine whether this is also the case in *ric8a* mutants, we next examined earlier-stage embryonic cortices. We found that at E14.5, similar to E16.5, there were already a large number of basement membrane breaches in the mutant cortex and almost all were associated with ectopic neurons (**Fig. S2**). By contrast, at E13.5, although there were breakages associated with ectopic neurons (**Fig. S2**), we in addition observed a significant number of sites with perturbed basement membrane but no associated ectopic neurons (**Fig. 3A-B”**). These results suggest that defects in the basement membrane likely precede ectopia formation and may a primary defect in *ric8a* mutants.

**Figure 3.**
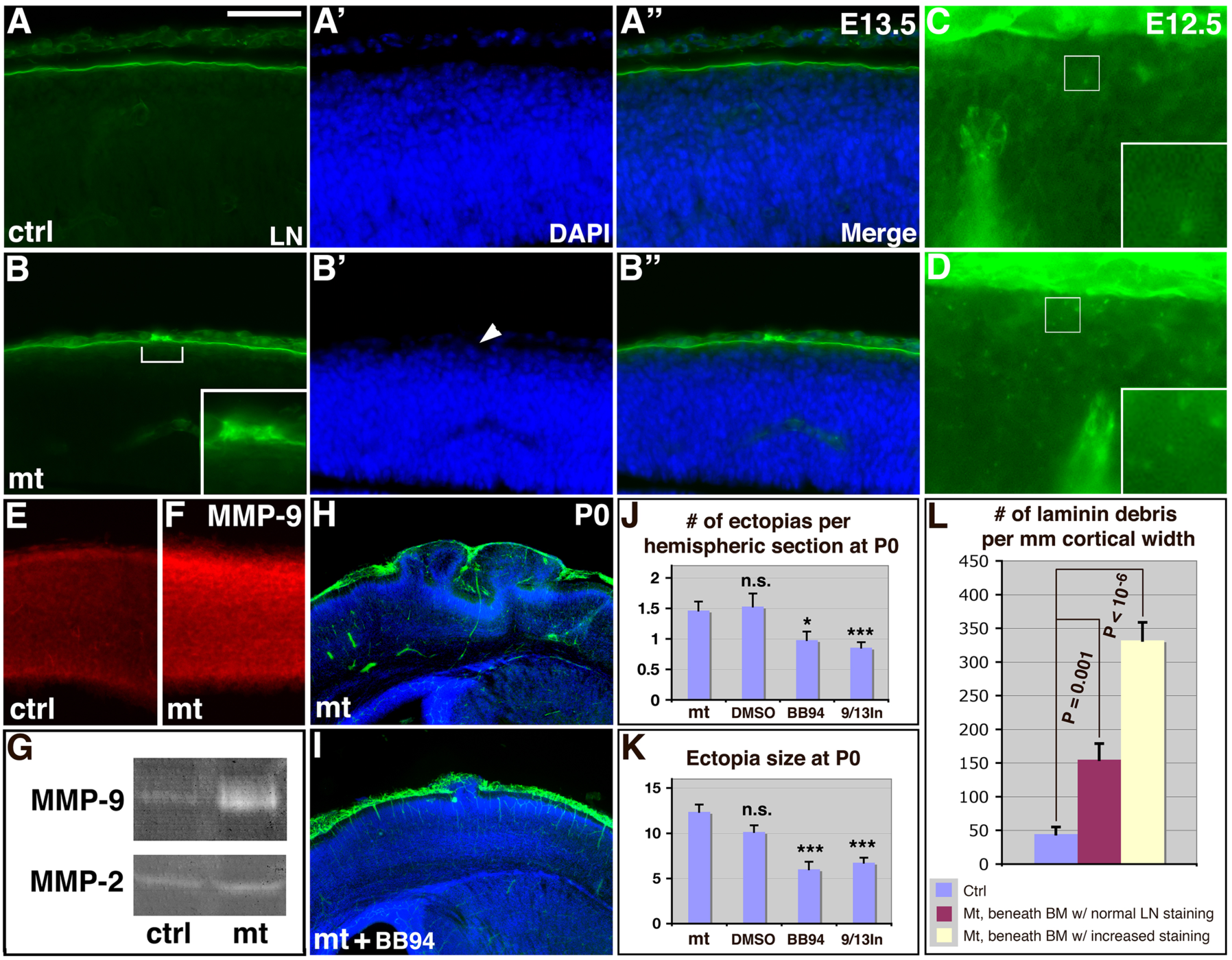
*ric8a* mutation results in elevated MMP9 levels, leading to pial basement membrane degradation. (**A**-**A”**) Laminin (LN, in green) and nuclear (DAPI, in blue) staining of control cortices at E13.5. (**B**-**B”**) Laminin and nuclear staining of *ric8a*/*emx1-cre* mutant cortices at E13.5. A small disruption of basement membrane is observed (bracket and inset in **B**), but not yet associated with ectopia (arrowhead in **B’**). (**C**, **D**, & **L**) Laminin (in green) staining of control (**C**) and mutant (**D**) cortices at E12.5. Increased numbers of laminin positive debris are observed in mutants (compare insets in **C** & **D**). Quantitative analysis shows significant increases (**L**). (**E** & **F**) MMP9 (in red) staining of control (**E**) and mutant cortices (**F**) at E13.5. Quantification shows statistically significant increases in mutants (see text). (**G**) Gel zymography of control and mutant cortical lysates at E13.5. Increased levels of MMP9 but not of MMP2 were observed in mutants. See quantification in text. (**H** & **I**) Laminin (in green) and nuclear (DAPI, in blue) staining of mutant cortices untreated (**H**) or treated (**I**) with BB94. (**J** & **K**) Quantitative analysis of ectopia number and size following MMP inhibitor treatment. *, *P* < 0.05; ***, *P* < 0.001; all compared to untreated mutants. Scale bar in (**A**), 100μm for (**A**-**B”**, **E** & **F**), 50μm for (**C** & **D)**, and 500μm for (**H** & **I**).

### Increased matrix metalloproteinases lead to basement membrane breach

As mentioned, in cobblestone lissencephaly, the most severe ectopia are observed at the midline (Beggs et al., 2003; Belvindrah et al., 2006; Graus-Porta et al., 2001; Huang et al., 2006; Moore et al., 2002; Niewmierzycka et al., 2005; Satz et al., 2010). In *ric8a* mutants, in contrast, we observed neither basement membrane breaches nor ectopias at the midline (**Fig. 1&2**). Radial glial fibers at ectopia sites, instead of retracting, also extend beyond the breach (**Fig. 2**). This suggests a distinct mechanism of basement membrane breach. Radial glial cells play an essential role in regulating cortical basement membrane integrity during development. To determine whether radial glial defects play a role in ectopia formation, we next examined whether radial glial cell fate is affected by *ric8a* mutation. We found that the cortical neural progenitor markers Pax6, Nestin, and Vimentin were all expressed normally in *ric8a* mutants (**Fig. S35A-H**). Radial glial proliferation was also normal (**Fig. S3I-N**). Furthermore, although *ric8a* has been found to regulate asymmetric cell division in invertebrates (Afshar et al., 2004; Couwenbergs et al., 2004; David et al., 2005; Hampoelz et al., 2005; Wang et al., 2005), we found no significant changes in the distribution of cleavage plane orientation of mitotic cells at the ventricular surface in *ric8a* mutants (**Fig. S3O-R**). These results indicate that radial glial defects are unlikely to be responsible for ectopia formation in *ric8a* mutants. Instead, at E12.5, while examining for potential basement membrane breaches (for which we found none), we observed increased numbers of laminin-positive debris in cortical regions immediately beneath the pia, both under basement membrane segments with normal laminin staining and segments with increased staining, the latter presumably marking future sites of breach (**Fig. 3C, D, L**). These results suggest that abnormal degradation of basement membrane may potentially be responsible for the formation of breaches and that ectopia activity of extracellular matrix proteinases might play a role.

To investigate this possibility, we evaluated the expression of several matrix metalloproteinases (MMPs) and found that immunoreactivity for MMP9 was increased in the mutants at E13.5 (**Fig. 3E, F**). Quantification showed that MMP9 staining intensity was significantly increased in the mutants, by ∼ 50% (control, 24.8 ± 0.2 AU (Arbitrary Units); mutant, 35.7 ± 1.7 AU; *P* = 0.002; n = 6). Similarly, gelatin gel zymography showed that the activity level of MMP9, but not that of MMP2, was also drastically increased in the mutants, at a level close to 4 times of that in controls (control, 1.00 ± 0.06 AU; mutant, 3.72 ±1.86 AU; *P* = 0.028; n = 4) (**Fig. 3G**). These results suggest that increased MMP activity may play a role in basement membrane breakdown and ectopia formation in ric8a mutants. To test this, we employed several MMP inhibitors to determine the effects of blocking MMP9 activity. First, we employed BB94, a broad-spectrum MMP inhibitor, to inactivate all MMPs. We found that daily BB94 administration from E12.5-14.5 significantly suppresses ectopia formation in *ric8a* mutants. The number of ectopia decreased by over 30% (mutant, 1.47 ± 0.14 / hemispheric section; mutant + BB94, 0.98 ± 0.14; *P* < 0.05, n = 50) and the size of ectopia decreased by over 50% (mutant, 12.37 ± 0.79 x 10^3^ μm^2^; mutant + BB94, 6.00 ± 0.86 x 10^3^ μm^2^; *P* < 0.001, n = 49) (**Fig. 3H-K**). By contrast, administration of DMSO alone had no significant effects on either the number (mutant, 1.47 ± 0.14 / hemispheric section; mutant + DMSO, 1.53 ± 0.21; *P* = 0.99, n = 51) or the size (mutant, 12.37 ± 0.79 x 10^3^ μm^2^; mutant + DMSO, 10.15 ± 0.73 x 10^3^ μm^2^; *P* = 0.24, n = 78) of the ectopia. These results indicate that MMP activity indeed functionally contributes to neuronal ectopia formation. We also employed an inhibitor specific for MMP9 and 13. We found that this inhibitor also significantly suppressed both the number, by over 40% (mutant, 1.47 ± 0.14 / hemispheric section; mutant + MMP9/13 inhibitor, 0.86 ± 0.09; *P* < 0.001, n = 90) and the size, by ∼45% (mutant, 12.37 ± 0.79 x 10^3^ μm^2^; mutant + DMSO, 6.75 ± 0.55 x 10^3^ μm^2^; *P* < 0.001, n = 77) of the ectopia (**Fig. 3J&K**). Together, these results demonstrate that increased MMP activity plays a key role in cortical basement membrane breach and ectopia formation in *ric8a* mutants.

### *ric8a* loss of function in non-neural cells causes ectopia

The increased MMP levels suggest potentially altered gene expression in cortices of *ric8a* mutants. To determine the specific cell type(s) responsible for ectopia formation, we employed a panel of cell type/developmental stage-specific *cre* lines for deleting *ric8a* (**Fig. 4**). As mentioned, using *nex-cre*, we found that *ric-8a* is not required in cortical plate neurons. To assess its role in marginal zone neurons, we employed *wnt3a-cre*, a *cre* line specifically expressed in the vast majority of Cajal-Retzius cells (Yoshida et al., 2006). Consistent with our interpretation against primary defects in Cajal-Retzius cells (Fig. 2), we found that *wnt3a-cre* mediated deletion also failed to perturb corticogenesis (**Fig. 4D-D”**). These results point to the involvement of cell types other than neurons. To further pinpoint the cell type(s) responsible, we next employed *nestin-cre*, a *cre* line targeting cortical neural progenitors beginning around E12.5 (Graus-Porta et al., 2001). Previous studies have shown that *emx1-cre* and *nestin-cre* target similar groups of cells involved in cortical basement membrane regulation as deletion of *β1 integrin* and related genes by either *cre* results in similar ectopia phenotypes (Belvindrah et al., 2006; Graus-Porta et al., 2001; Huang et al., 2006; Niewmierzycka et al., 2005). To our surprise, unlike *emx1-cre* **(Fig. 1)**, we found that deletion of *ric8a* by *nestin-cre* did not result in any basement membrane breach or ectopia (**Fig. 4C-C”**). This suggests that *ric8a* is likely not required in cortical neural progenitors (radial glial cells) for maintaining basement membrane integrity. Since *nestin-cre*-mediated deletion in neural progenitors is also inherited by cortical neurons and glia, these results also indicate that the combined function of *ric8a* in these neural cell types is not required for maintaining basement membrane integrity. The onset of *nestin-cre* expression, however, is slightly delayed than *emx1-cre* (Gorski et al., 2002). To definitely exclude a role played by this temporal difference, we employed *foxg1-cre*, a *cre* line expressed in neural progenitors from the very onset of forebrain development (Hebert and McConnell, 2000). We found that *foxg1-cre* deletion of *ric8a* also failed to result in cortical ectopia (**Fig. 4E-E”**). Together, these results rule out potential cell-autonomous defects in neural lineage cells as responsible for basement membrane degradation and ectopia formation in *ric8a* mutants and suggest a role played by non-neural cell types.

**Figure 4.**
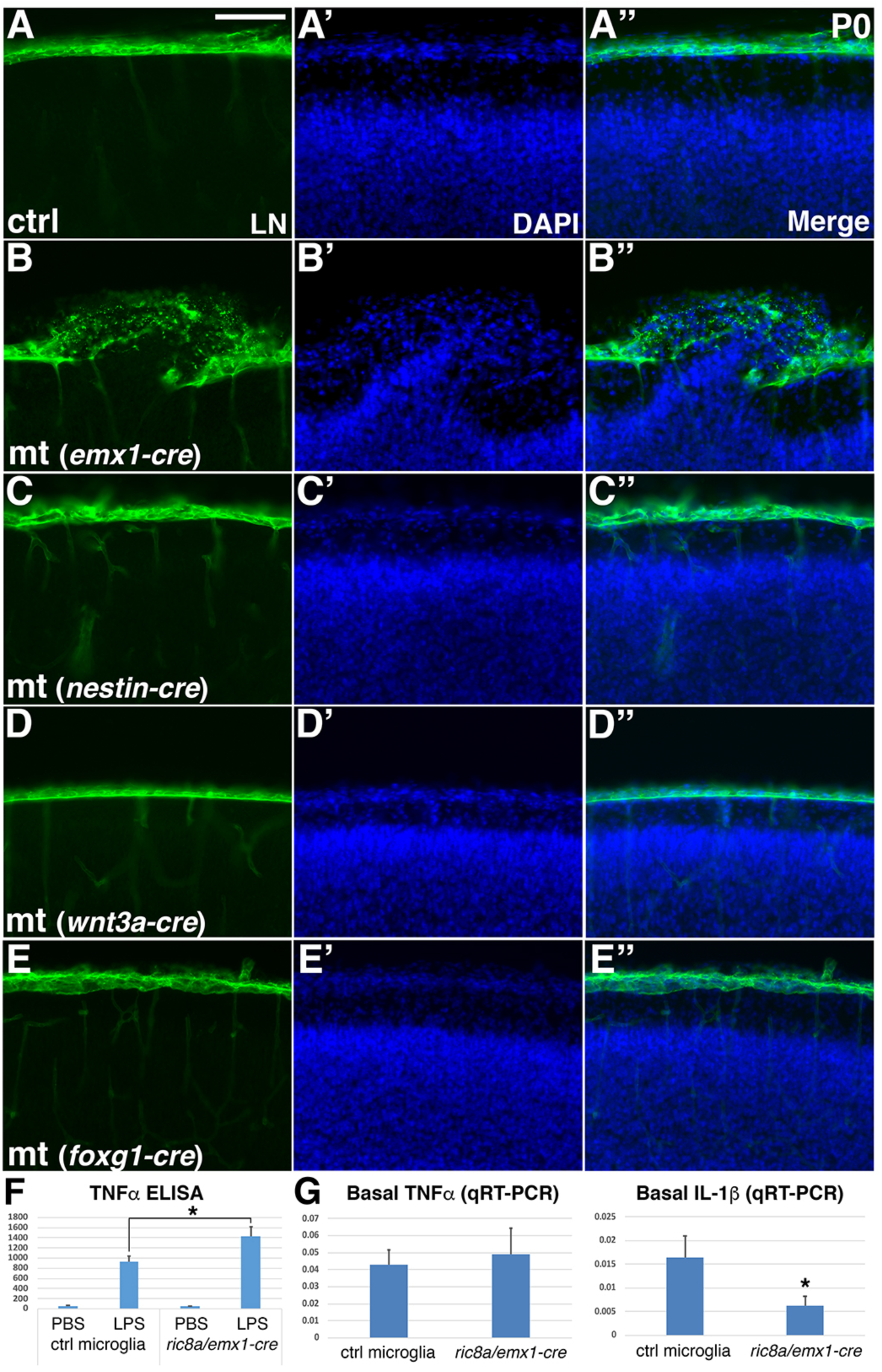
*ric8a* deficiency in neural lineage cells does not account for ectopia formation. (**A-A”**) Laminin (LN, in green) and nuclear (DAPI, in blue) staining of control cortices at P0. A continuous basement membrane is observed at the pia, beneath which cells are well organized in the cortical wall. (**B**-**B”**) Staining of *ric8a*/*emx1-cre* mutant cortices at P0. Basement membrane breach and neuronal ectopia are observed following *ric8a* deletion by *emx1-cre*, a *cre* line expressed in cortical radial glial progenitors beginning at E10.5. **(C**-**C”**) Staining of *ric8a*/*nestin-cre* mutant cortices at P0. No obvious basement membrane breach or neuronal ectopia is observed following *ric8a* deletion by *nestin-cre*, a *cre* line expressed in cortical progenitors beginning around E12.5. **(D**-**D”**) Staining of *ric8a*/*wnt3a-cre* mutant cortices at P0. No obvious basement membrane breach or neuronal ectopia is observed following *ric8a* deletion by *wnt3a-cre*, a *cre* line expressed in Cajal-Retzius cells. (**E-E”**) Staining of *ric8a*/*foxg1-cre* mutant cortices at P0. No obvious basement membrane breach or neuronal ectopia is observed following *ric8a* deletion by *foxg1-cre*, a *cre* line expressed in forebrain neural progenitors from E9.0. (**F-G**) TNFα secretion (pg/ml) (**F**) and basal TNFα and IL-1β mRNA expression (**G**) in control and *ric8a/emx1-cre* mutant microglia. *, *P* < 0.05; n = 5-8 each group. Scale bar in (**A**), 100μm for (**A-E”**).

During early embryogenesis, much of the neural tube at the dorsal midline (except in the forebrain) undergoes a process similar to epithelial-mesenchymal transition that gives rises to neural crest cells. This process, regulated by diffusible factors including Wnt, involves a process of targeted basement membrane breakdown similar to what we observed in *ric8a* mutants. To determine whether ectopic epithelial-mesenchymal transition may potentially play a role in the *ric8a* ectopia phenotype, we also examined the expression of a number of neural crest development markers. We found that none of the canonical neural crest markers including Msx1/2, Pax3, and NTFRp75 were ectopically expressed in *ric8a/emx1-cre* mutant cortices (data not shown). There were also no obvious changes in the level of Wnt pathway activation in the mutant cortex (**Fig. S4A-D**). Furthermore, neither the expression pattern nor the density of cells positive for the layer-specific markers Cux1 and Ctip2 were affected in most cortical areas (**Fig. S4E-K**). These results suggest that ectopic activation of an epithelial-mesenchymal transition related process is unlikely to play a role in ectopia formation in *ric8a* mutants.

### *ric8a* deficiency in microglia results in cortical ectopia formation

Having ruled out roles of *ric8a* deficiency in neural lineage cells, we next focus on non-neural cell types in the brain. The cerebral cortex contains two main non-neural cell populations, microglia and endothelial cells. To determine which cell type is responsible, we re-examined *emx1* gene expression. Using an RNA-seq database that systematically analyzes all cortical cell type gene expression (Zhang et al., 2014), we found that *emx1* is expressed at a significant level in microglial but not in endothelial cells. To test whether *emx1-cre* is similarly expressed as the endogenous gene and active in this cell type, we cultured microglia from *ric8a/emx1-cre* mutant cortices. We found that relative to wildtype cells, mutant microglia showed significantly elevated secretion of tumor necrosis factor α (TNFα) (**Fig. 4F**). At the transcriptional level, basal expression of interleukin-1β (IL-1β) is also altered (**Fig. 4G**). These results indicate that *emx1-cre* is indeed active in microglia.

To further determine the effects of microglial *ric8a* deficiency, we next employed a microglia-specific *cx3cr1-cre* (Yona et al., 2013). Similar to those from *ric8a/emx1-cre* mutants, microglia from *ric8a/cx3cr1-cre* mutants also showed strongly elevated secretion of a large number of cytokines including TNFα, IL-1β, and IL-6 upon stimulation by lipopolysaccharide (LPS) (**Fig. 5A**). At the transcriptional level, IL-1β basal expression is also altered in comparison to that in wildtype cells (**Fig. 5B**). Furthermore, all these cytokines showed elevated transcriptional expression in mutant microglia upon stimulation by LPS (**Fig. 5B**). Similar elevated responses were also obseved in mutant microglia upon immune stimulation with polyinosinic-polycytidylic acid (poly I:C), which activates different TLR receptors (data not shown). Thus, these results indicate that *ric8a* deficiency results in a general microglial hypersensitivity to immune stimulation.

**Figure 5.**
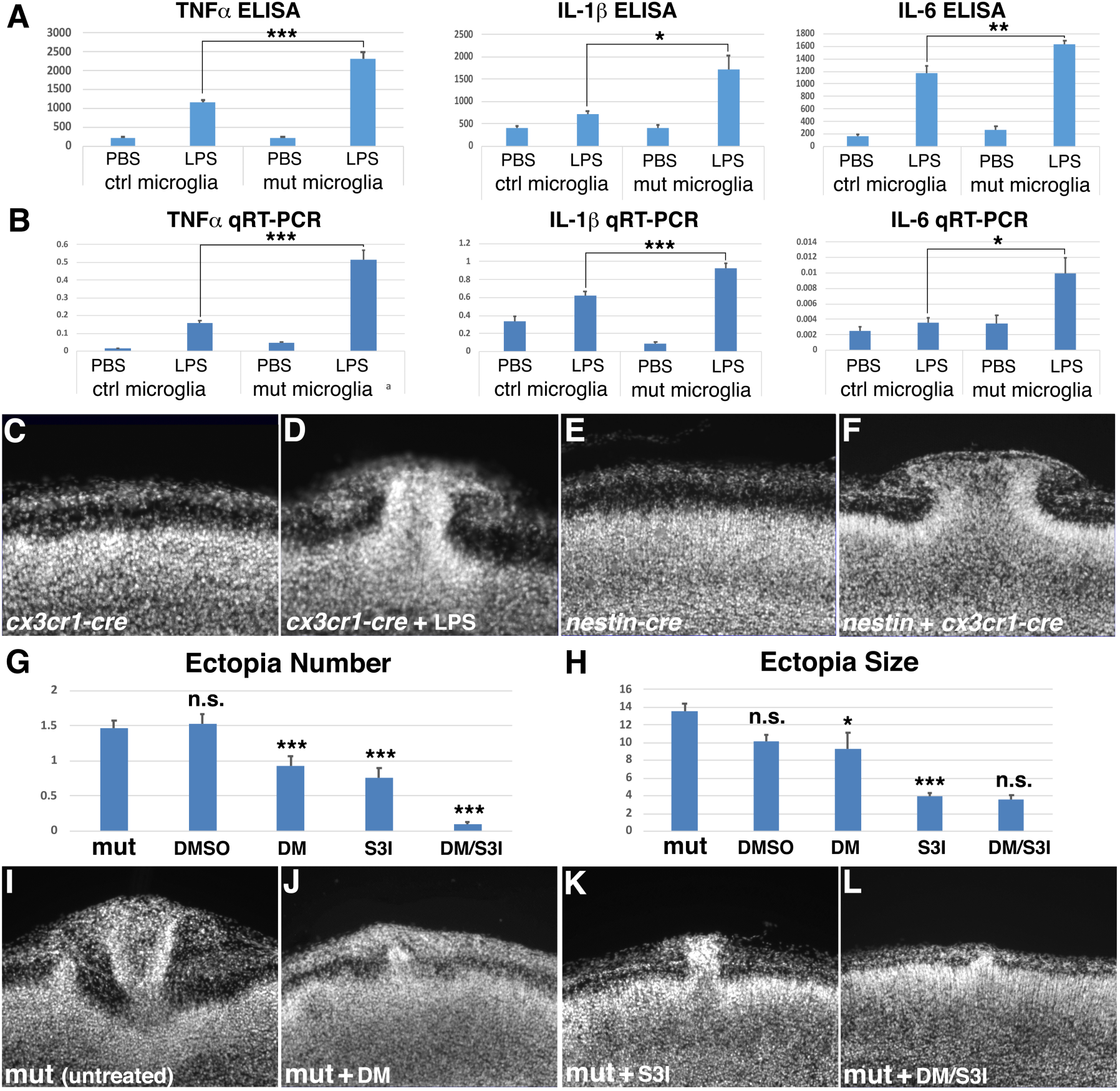
*ric8a* deficiency in microglia results in cortical ectopia. (**A**) TNFα, IL-1β, and IL-6 secretion (pg/ml) in control and *ric8a/cx3cr1-cre* mutant microglia following LPS stimulation. *, *P* < 0.05; **, *P* < 0.01; ***, *P* < 0.001; n = 6-8 each group. (**B**) TNFα, IL-1 β, and IL-6 mRNA expression in control and *ric8a/cx3cr1-cre* mutant microglia following LPS stimulation. *, *P* < 0.05; ***, *P* < 0.001; n = 5-6 each group. (**C**-**D**) Nuclear (DAPI, in grey) staining of *ric8a/cx3cr1-cre* mutant cortices at P0 in the absence (**C**) or presence (**D**) of LPS treatment during embryogenesis. (**E-F**) Nuclear (DAPI, in grey) staining of *ric8a/nestin-cre* single *cre* (**E**) and *ric8a/nestin-cre+cx3cr1-cre* double *cre* (**F**) mutant cortices at P0. **(G-H**) Quantitative analysis of ectopia number (**G**) and size (**H**) in the neonatal mutant cortex after DMSO, DM, S3I, and DM/S3I dual treatment at E12.5. *, *P* < 0.05; ***, *P* < 0.001; all compared to untreated mutants. The reduction in ectopia size after dual treatment is not statistically significant, likely because of the small number of ectopias that remained. **(I**-**L**) Nuclear (DAPI, in grey) staining of untreated (**I**), DM (**J**), S3I (**K**), and DM/S3I (**L**) dual treated mutant cortices at P0. Scale bar in (**C**), 100 μm for (**C**-**F**) and 150 μm for (**I-L**).

To determine whether *ric8a* deficiency in microglia is sufficient to cause cortical ectopia, we next examined *ric8a/cx3cr1-cre* mutant cortices. We found that cortical layering is normal without ectopia in these animals (**Fig. 5C**). This is consistent with our observation that *ric8a* mutant microglia have largely normal activity under basal conditions in vitro and only show elevated cytokine release relative to wildtype cells upon immune stimulation (**Fig. 5A-B**). To mimic immune stimulation, we administered LPS to pregnant mothers of mutant animals at E11.5-12.5. While no cortical ectopia were observed in any of the control littermates treated with LPS (32 of 32 examined), we observed ectopia in over 50% of *ric8a/cx3cr1-cre* mutant neonates (10 of 19 mutant neonates examined) (**Fig. 5D**). In *emx1-cre* mutants, we observe cortical ectopia in the absence of LPS administration (**Fig. 1**). We suspected that this may be due to concurrent *ric8a* deficiency in cortical neural cells in these mutants, which may result in neural tissue damages that serve as immune stimulants for the mutant microglia. To test this, we removed *ric8a* from neural cells in *cx3cr1-cre* mutants by additionally introducing *nestin-cre*. As mentioned, *ric8a* deletion by *nestin-cre* alone does not result in cortical ectopia (**Fig. 4C**). However, we found that the combined deletion of *ric8a* by *cx3cr1-cre* and *nestin-cre* resulted in cortical ectopia formation in all of the double *cre* mutant animals (6 of 6 neonates examined) (**Fig. 5F**). Thus, these results indicate that *ric8a* deficiency in microglia ultimately leads to cortical ectopia formation in *ric8a* mutants.

### Inhibition of immune activation suppresses ectopia formation

If inappropriate microglial activation is responsible for ectopia formation in *ric8a* mutants, inhibition of immune response should suppress the phenotype. To test this, we employed dorsomorphin (DM) and S3I-201(S3I), which are inhibitors of AMPK and Stat3 activity as well as inflammatory signaling (Covarrubias et al., 2016; Qin et al., 2012). We found that administration of dorsomorphin during early corticogenesis significantly suppresses ectopia formation (**Fig. 5G, H, J**), reducing both the number, by close to 40% (mutant, 1.47 ± 0.11 / hemispheric section; mutant + dorsomorphin, 0.92 ± 0.14; *P* = 0.003; n = 51) and the size, by over 30% (mutant, 13.53 ± 0.88 x 10^3^ μm^2^; mutant + dorsomorphin, 9.28 ± 1.82 x 10^3^ μm^2^; *P* = 0.045, n = 47) of the ectopia. Similarly, administration of S3I-201 also significantly suppressed ectopia formation (**Fig. 5G, H, K**), reducing both the number, by close to 50% (mutant, 1.47 ± 0.11 / hemispheric section; mutant + S3I-201, 0.77 ± 0.13; *P* < 0.001; n = 43) and the size, by over 70% (mutant, 13.53 ± 0.88 x 10^3^ μm^2^; mutant + S3I-201, 3.91 ± 0.39 x 10^3^ μm^2^; *P* < 0.001, n = 33) of the ectopia. By contrast, treatment with DMSO alone had no significant effects on either ectopia number (mutant, 1.47 ± 0.11 / hemispheric section; mutant + DMSO, 1.53 ± 0.13; *P* = 1.00; n = 51) or ectopia size (mutant, 13.53 ± 0.88 x 10^3^ μm^2^; mutant + DMSO, 10.15 ± 0.73 x 10^3^ μm^2^; *P* = 0.107, n = 78) (**Fig. 5G-H**). Most importantly, we found that simultaneous administration of both dorsomorphin and S3I-201 eliminates almost all ectopia, decreasing the number by close to 94% (mutant, 1.47 ± 0.11 / hemispheric section; mutant + dorsomorphin/S3I-201, 0.09 ± 0.04; *P* < 0.001; n = 54) (**Fig. 5G, L**). Thus, these results further demonstrate that inappropriate microglial inflammatory activation plays a key role in ectopia formation in *ric8a* mutants.

Activated macrophages induces extracellular matrix proteinase expression during immune response to facilitate migration (Hanania et al., 2012). We observed elevated levels of MMP9 in *ric8a* mutant cortices (**Fig. 3E-G**). We also found inhibition of MMPs significantly suppress ectopia formation in ric8a mutants (**Fig. 3H-K**). These results suggest that inappropriate microglial activation in *ric8a* mutants may lead to ectopia formation through increased proteinase expression. To test if this is the link, we next examined effects of immune suppression on MMP9 activity in *ric8a/emx1-cre* cortices. We found that treatment with dorsomorphin and S3I-201 restored levels of MMP9 in the mutant cortices to levels comparable to those in control animals (**Fig. S5A-D**). Microglial activation also leads to astrogliosis, a secondary response by astrocytes. Consistent with the suppression of microglial activation, we found that the staining intensity of GFAP, a marker for astrogliosis, was also significantly down-regulated in *ric8a/emx1-cre* mutant cortices following dorsomorphin and S3I-201 treatment (**Fig. S5E-J**). Thus, these results further support the interpretation that *ric8a* deficiency in microglia results in cortical ectopia through elevating cortical MMP9 levels.

### *app* mutation in microglial cells results in similar ectopia phenotype

As mentioned, a unique feature of the ectopia observed in *ric8a* mutants is their exclusive localization to the lateral cortex. Only a few other mutants show this feature. One of them is the triple mutant in *app* family genes *app/aplp1/2* (Herms et al., 2004). Another is a related double mutant in *apbb1/2*, which encode a family of adaptors binding to the cytoplasmic domain of APP (Guenette et al., 2006). The similarity of these phenotypes raises the possibility that inappropriate microglial activation may also be responsible for ectopia formation in these *app*-related mutants. Consistent with this, *app* knockdown in cortical neurons has been found to result in under-instead of over-migration (ectopia) of cortical neurons (Young-Pearse et al., 2007). To test this, we first measured the activity of microglia from *app/cx3cr1-cre* microglial specific conditional mutants (Wang et al., 2009). We found that TNFα and IL-6 secretion by *app/cx3cr1-cre* mutant microglia was significantly altered (**Fig. 6A, S6A**). At the transcriptional level, induction of IL-6 mRNA was also muted (**Fig. S6B**). These results indicate that *app* indeed normally regulates microglial activity in a cell-autonomous manner.

**Figure 6.**
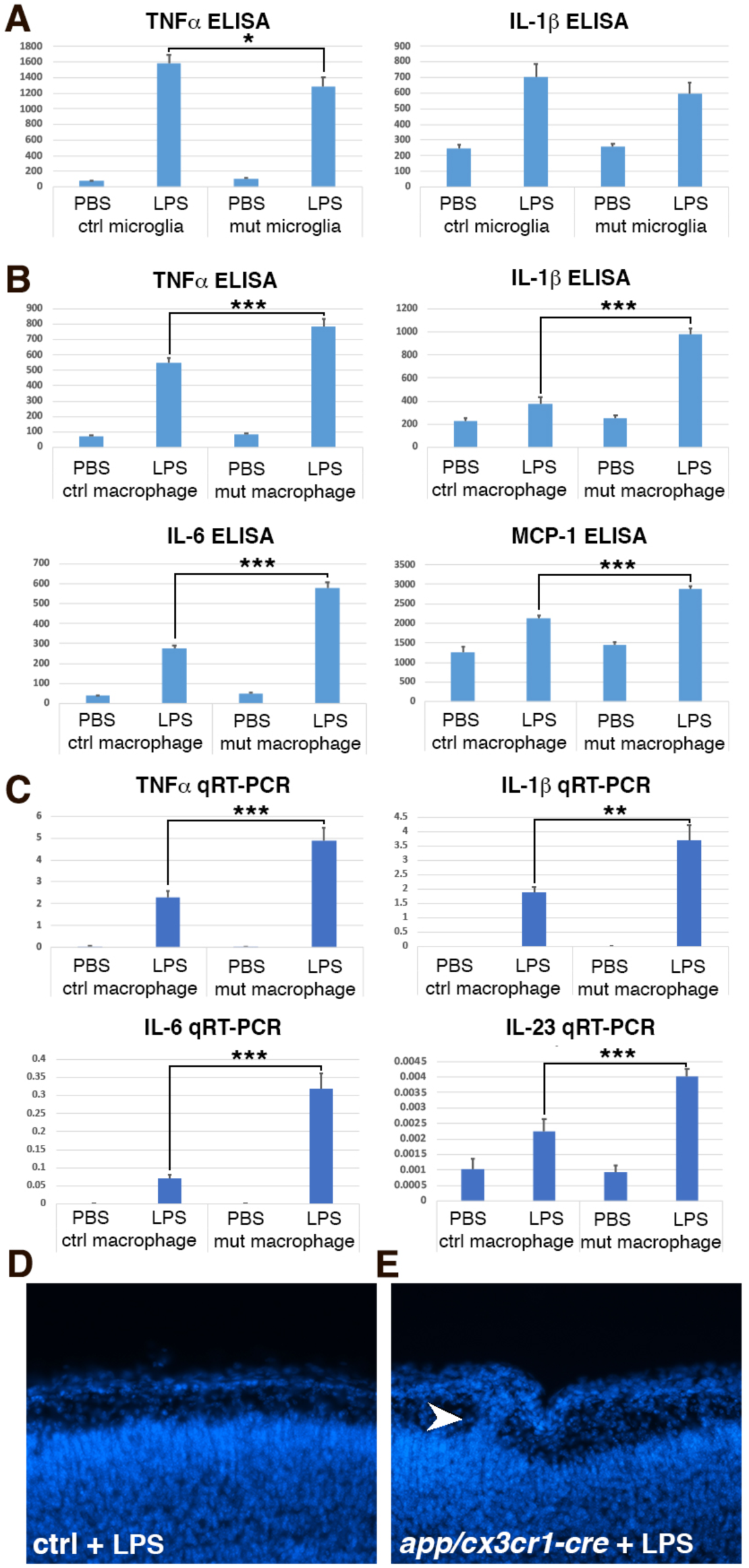
*app* mutation results in microglial lineage cell hypersensitivity in vitro and cortical ectopia in vivo. **(A)** TNFα and IL-1β secretion (pg/ml) in control and *app/cx3cr1-cre* mutant microglia following LPS stimulation. *, *P* < 0.05; n = 7-9 each group. (**B**) TNFα, IL-1β, IL-6, and MCP1 secretion (pg/ml) in fresh unelicited control and *app/cx3cr1-cre* mutant peritoneal macrophages following LPS stimulation. ***, *P* < 0.001; n = 7-10 each group. (**C**) TNFα, IL-1 β, IL-6, and IL-23 mRNA expression in fresh unelicited control and *app/cx3cr1-cre* mutant peritoneal macrophages following LPS stimulation. **, *P* < 0.01; ***, *P* < 0.001; n = 6 each group (**D** & **E**) Nuclear (DAPI, in blue) staining of control (**D**) and LPS-treated *app/cx3cr1-cre* mutant (**E**) cortices at P0. Note cortical ectopia in the mutant cortex (arrowhead). Scale bar in (**D**), 200μm for (**D-E**).

Microglia show attenuated immune response following prolonged exposure to stimulants, especially when harboring strong loss-of-function mutations in inflammation-regulating pathways (Chamberlain et al., 2015; Sayed et al., 2018). We suspect that the attenuated cytokine expression by *app* mutant microglia may result from similar effects. To bypass this complication, we isolated fresh, unelicited peritoneal macrophages from *app/cx3cr1-cre* mutant animals and directly determined their activity without culturing. We found that *app* mutant macrophages showed strongly elevated secretion of all cytokines tested, including TNFα, IL-1β, IL-6, and MCP-1, upon immune stimulation (**Fig. 6B**). At the transcriptional level, the induction of TNFα, IL-1β, IL-6, and IL-13a mRNAs was also all strongly increased compared to that of control cells (**Fig. 6C**). These results indicate that, like *ric8a*, *app* gene also normally negatively regulates immune activation by microglial lineage cells. Since the activation of *ric8a* mutant microglia in vivo results in cortical ectopia formation, we next determined whether activating *app* mutant microglia had similar effects. To this end, we administered LPS to *app/cx3cr1-cre* mutant animals during embryogenesis, as we did to *ric8a/cx3cr1-cre* mutants. We found that a significant number of *app/cx3cr1-cre* mutants indeed showed cortical ectopia after LPS treatment (6 of 31 neonates examined, ∼19%), while none of the LPS-treated littermate controls showed ectopia (0 of 81 neonates examined) (**Fig. 6E**). The severity of the ectopia, however, appeared to be lower than that observed in *ric8a/cx3cr1-cre* mutants. This is likely in part a result of the reduced LPS dosage (by ∼3 folds) we had to use in these animas due to the enhanced immune sensitivity of the *app* genetic background. Other *app* gene family members are also expressed in microglia (Zhang et al., 2014) and may also functionally compensate. Together, these results indicate that *app* also negatively regulates microglial activity in a cell-autonomous manner in vivo. The similarities of *app* to *ric8a* mutant phenotypes, both in vitro and in vivo, further suggest that they may form a pathway in this process.

### Aβ monomers suppress microglial activity via an APP and Ric8a-dependent pathway

Our results so far indicate that APP and Ric8a may form a signaling pathway in microglia that negatively regulates microglial activation in the brain. This raises questions on the identity of the endogenous ligands that may regulate this pathway. Previous studies have reported several diffusible molecules that bind to APP and/or activate APP-dependent pathways (Fogel et al., 2014; Milosch et al., 2014; Rice et al., 2012). Among them, the APP cleavage product Aβ has been shown to bind to APP with picomolar sensitivity (Fogel et al., 2014; Shaked et al., 2006). In the brain, Aβ is secreted mainly by neurons and its production is increased upon neural cell exposure to cytokines (Blasko et al., 1999; Buxbaum et al., 1992; Del Bo et al., 1995; He et al., 2007; Liao et al., 2004; Ohyagi and Tabira, 1993; Rogers et al., 1999). Since cytokines are mainly produced by microglia and other glia in the brain, this raises the possibility that Aβ may potentially serve as a neuron-derived negative feedback signal that helps maintain brain immune homeostasis. Aβ peptides exist in forms including Aβ40 and Aβ42. When diluted from DMSO into aqueous solutions, Aβ42 peptides readily form oligomeric structures, while Aβ40 peptides do not (LeVine, 2004; Stine et al., 2011). Studies have reported pro-inflammatory effects of Aβ fibrils on microglia by themselves (Bamberger and Landreth, 2001; Bianca et al., 1999; Casal et al., 2004; Combs et al., 2001; Glass et al., 2010; Halle et al., 2008; Ishii et al., 2000; Lee et al., 1993; Lorton et al., 1996; McDonald et al., 1997; Muehlhauser et al., 2001; Reed-Geaghan et al., 2009; Richard et al., 2008; Tan et al., 1999; Van Muiswinkel et al., 1999; Weldon et al., 1998). Non-fibrillar Aβ, in contrast, has been found to lack pro-inflammatory effects by itself (Halle et al., 2008; Lorton et al., 1996; Muehlhauser et al., 2001). However, these experiments have done by using non-fibrillar Aβ alone and the effects of non-fibrillar Aβ on microglial activation by immune stimuli have not been tested. In fact, in the periphery, both Aβ40 and Aβ42, when prepared in conditions favoring the monomer conformation, have been found to inhibit T cell activation (Grant et al., 2012). This raises the question of whether Aβ monomers may also have inhibitory activity against microglial activation in the brain and if so, by what mechanism.

To test the effects of Aβ monomers on microglial activation, we prepared Aβ40 stock solutions using DMSO, which preserves Aβ40 predominantly in the monomeric conformation (LeVine, 2004; Stine et al., 2011), and treated microglia with Aβ40 to determine the effects on microglial activation by different immune stimuli. We found that Aβ40 monomers potently suppressed LPS-induced secretion of a large number of cytokines including TNFα, IL-6, and MCP-1 as well as LPS-primed ATP-induced secretion of IL-1β, by up to 40% (**Fig. 7A**). These effects were observed at concentrations as low as 50 nM (**Fig. S7A**). Similarly, we also found that Aβ40 monomers potently suppressed poly I:C-induced cytokine secretion (**Fig. 7B**). Furthermore, these effects were consistently observed with Aβ40 peptides obtained from different commercial sources (**Fig. S7B**). At the transcriptional level, Aβ40 monomers also potently suppress the induction of cytokines including IL-1β IL-6, IL-10, and IL-23a, by up to 60% (**Fig. 7C, S7C**). Thus, these results demonstrate that monomeric Aβ has a strong anti-inflammatory activity that negatively regulates microglial activation by different immune stimuli.

**Figure 7.**
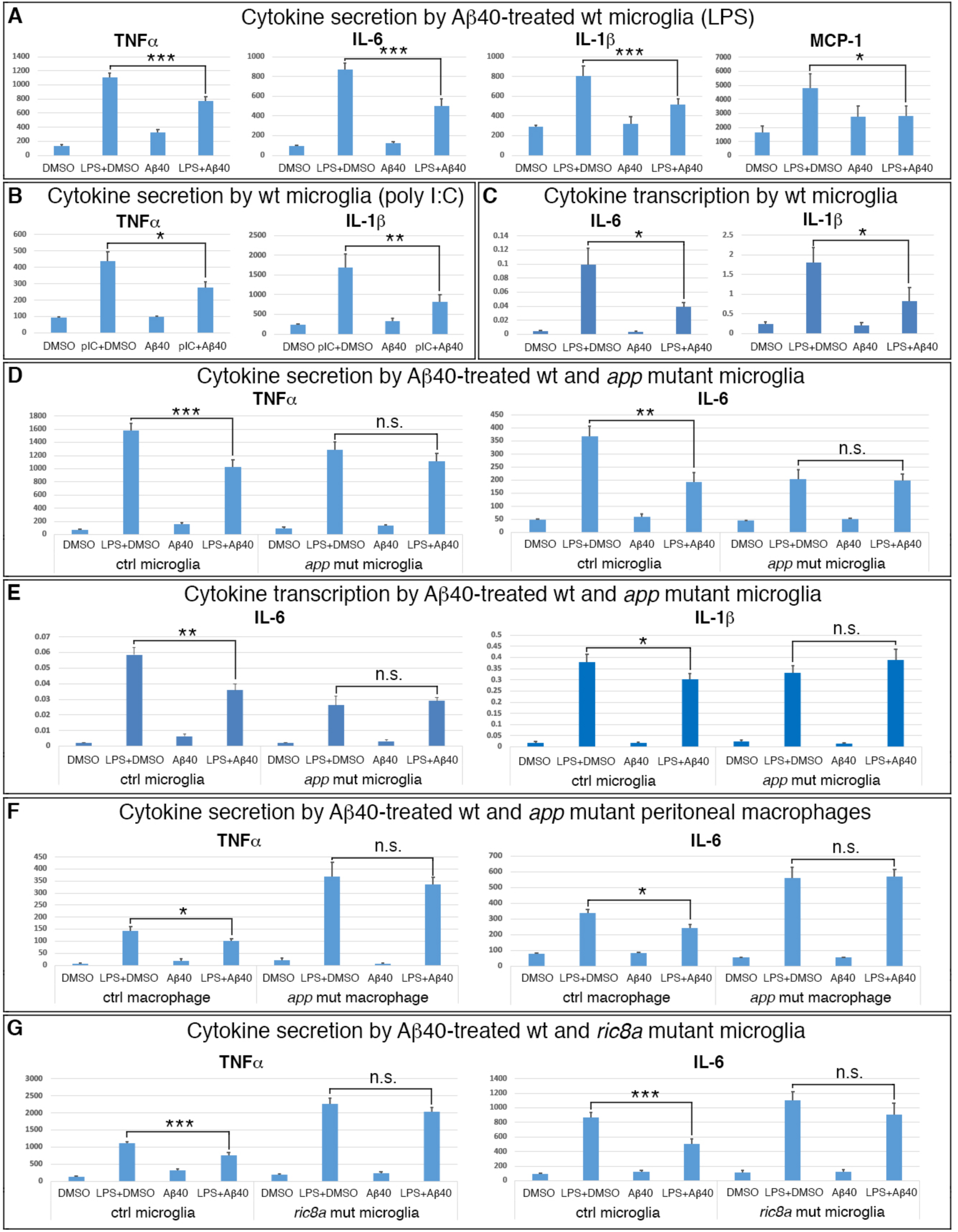
Aβ monomers suppress microglial immune activation via an APP and Ric8a-dependent pathway. **(A)** TNFα, IL-6, IL-1β, and MCP1 secretion (pg/ml) by wildtype microglia following LPS stimulation in the absence or presence of Aβ40 (200 or 500nM). *, *P* < 0.05; ***, *P* < 0.001; n = 8-14 each group. **(B)** TNFα and IL-1β secretion (pg/ml) by wildtype microglia following poly I:C stimulation in the absence or presence of Aβ40 (500nM). *, *P* < 0.05; **, *P* < 0.01; n = 6-7 each group. **(C)** IL-6 and IL-1β mRNA induction in wildtype microglia following LPS stimulation in the absence or presence of Aβ40 (500nM). *, *P* < 0.05; n = 6 each group. **(D)** TNFα and IL-6 secretion (pg/ml) by control and *app/cx3cr1-cre* mutant microglia following LPS stimulation in the absence or presence of Aβ40 (200nM). **, *P* < 0.01; ***, *P* < 0.001; n = 8 each group. **(E)** IL-6 and IL-1β mRNA induction in control and *app/cx3cr1-cre* mutant microglia following LPS stimulation in the absence or presence of Aβ40 (200nM). *, *P* < 0.05; **, *P* < 0.01; n = 6 each group. **(F)** TNFα and IL-6 secretion (pg/ml) by control and *app/cx3cr1-cre* mutant peritoneal macrophages following LPS stimulation in the absence or presence of Aβ40 (500nM). *,*P* < 0.05; n = 6-7 each group **(G)** TNFα and IL-6 secretion (pg/ml) by control and *ric8a/cx3cr1-cre* mutant microglia following LPS stimulation in the absence or presence of Aβ40 (200nM). ***, *P* < 0.001; n = 12-14 each group.

To determine whether Aβ monomers signals through APP in the suppression of microglial activation, we next employed *app* mutant cells. While Aβ40 monomers potently suppressed cytokine secretion by microglia derived from control littermates, we found it fail to suppress cytokine secretion by microglia derived from *app/cx3cr1-cre* mutants, including that of TNFα, IL-6, and IL-1β (**Fig. 7D, S7D**). These effects are specific to *app* since Aβ40 still significantly suppressed cytokine secretion by *aplp2* mutant microglia (**Fig. S7E**). At the transcriptional level, Aβ40 monomers also potently suppressed the transcriptional induction of IL-1β, IL-6, IL-10, and IL-23 in control, but not in *app/cx3cr1-cre* mutant microglia (**Fig. 7E, S7F**). Together, these results indicate that Aβ monomers signal through an APP-dependent microglial pathway to suppress microglial immune activation. Microglia derived from *app/cx3cr1-*cre mutants show attenuated immune activation compared to wildtype cells. To assess whether this may affect microglial response to Aβ monomers, we also tested fresh *app/cx3cr1-cre* mutant macrophages. We found that, even though *app/cx3cr1-cre* mutant macrophages showed elevated response to immune stimulation, they still failed to respond to Aβ40 and instead displayed levels of cytokine secretion that are statistically identical to those by DMSO-treated cells (**Fig. 7F, S7G**). These results further strengthen the conclusion that Aβ40 signals through an APP-dependent pathway to suppress microglial activation.

To determine whether Ric8a plays a role in mediating Aβ monomer effects on microglial activation, we next employed *ric8a* mutant cells. Similar to *app* cells, we found that, while Aβ40 monomers potently suppressed TNFα and IL-6 secretion by microglia derived from control littermates, they failed to suppress the secretion of either TNFα or IL-6 by *ric8a/cx3cr1-cre* mutant microglia (**Fig. 7G**). These results indicate that Aβ monomer suppression of TNFα and IL-6 secretion also depends on Ric8a, and thus heterotrimeric G protein, function in microglia. Interestingly, in contrast to TNFα and IL-6, we found that Aβ40 still potently suppressed IL-1β secretion by *ric8a/cx3cr1-cre* mutant microglia (**Fig. S7H**). Furthermore, the transcriptional induction of IL-6 mRNA in *ric8a* mutant cells is also still significantly suppressed by Aβ40 (**Fig. S7I**). These results indicate that while some of the anti-inflammatory effects of Aβ monomers are mediated by Ric8a-dependent pathways, others appear to be independent. Altogether, these results uncover a novel Aβ monomer-activated APP/Ric8a-mediated anti-inflammatory pathway in microglia, the perturbation of which results in neuronal ectopia and laminar disruption in cortical development.

## DISCUSSION

Microglia, one of the few non-neural cell types of the brain, originate from outside the nervous system and populate the brain during early development. They play crucial roles in the development, function, and plasticity of the brain neural circuitry and are critically involved in many diseases, especially those associated with aging and neurodegeneration. Because microglia are immune cells that express large numbers of cytokines that can cause highly deleterious effects if uncontrolled, microglial homeostasis is critical to the health of the brain. In this article, we uncover a previously unknown anti-inflammatory signaling pathway in microglia that regulates microglial homeostasis. We find that disruption of this pathway results in abnormal microglial activation during early cortical development, leading to elevated matrix proteinase activity, cortical basement membrane breakdown, and neuronal ectopia formation. We show that signaling by this pathway depends on the function of the transmembrane protein APP as well as Ric8a-regulated heterotrimeric G proteins. Furthermore, we show that monomers of the APP cleavage product αβ potently suppress microglial cytokine transcription and secretion through this pathway. These results provide new insights into the biological activity of Aβ as well as the intercellular communication mechanisms that regulate microglial homeostasis in the brain. They may facilitate better understanding of the pathogenic mechanisms underlying Alzheimer’s disease.

### Microglial activity and cortical ectopia

The pial basement membrane of the cerebral cortex plays a critical role in the normal development of the brain and the assembly of the neural circuitry. In cobblestone lissencephaly, genetic defects in the dystroglycan cell adhesion complex result in compromised interactions between radial glia and basement membrane, leading to basement membrane breach, neuronal ectopia formation, defective cortical lamination, and perturbed brain function (Moore et al., 2002; Satz et al., 2010; Waite et al., 2012). Similarly, deficiency in the integrin adhesion complex, which cooperates with the dystroglycan complex in the assembly and maintenance of the cortical basement membrane, also results in pial neuronal ectopia and cortical lamination defects (Beggs et al., 2003; Belvindrah et al., 2006; Graus-Porta et al., 2001; Huang et al., 2006; Jeong et al., 2013; Li et al., 2008; Niewmierzycka et al., 2005). While this body of studies has convincingly implicated defective basement membrane assembly and maintenance as a major factor in the development of cobblestone lissencephaly, it remains unclear whether all types of basement membrane breaches are of the same etiology. In *ric8a* mutants, we find that excessive microglial activity appears to be a primary cause of basement membrane breach, while radial glial deficiency appears to play no significant role (**Figs. 3-5**) Specifically, we find that abnormal microglial activation leads to increased MMP activity in *ric8a* mutants, causes basement membrane breach, and results in ectopia formation. These results indicate that morphologically similar ectopia in the brain may arise through different etiologies. We also find that administration of anti-inflammatory drugs can almost eliminate ectopia caused by microglial overactivation (**Fig. 5**), while administration of MMP inhibitors also suppresses the ectopia (**Fig. 3**). These results provide new insights into the cause and potential prevention of cobblestone lissencephaly related brain disorders.

### Aβ, APP, and brain immune homeostasis

Genetic mutations in APP are well known to be associated with the development of familial Alzheimer’s disease in humans. The APP cleavage product Aβ accumulates in large quantities in plaques in the Alzheimer’s disease brain and plays critical roles in the disease pathology. However, the normal biological functions of APP and of Aβ in the brain still remain largely obscure. APP belongs to a gene family with three members that show generally subtle defects when mutated individually in mice (Muller and Zheng, 2012). Only triple mutations in the *app* family genes result in a severe phenotype of cortical ectopia (Herms et al., 2004). Our results show that, like those in *ric8*a mutants, these ectopia likely result from uncontrolled microglial activity during cortical development (**Fig. 6**). We find *app* mutant microglial lineage cells show hypersensitivity to immune stimulation. We also find that, when stimulated, microglial specific *app* deletion results in cortical ectopia similar to those *in ric8a* as well as *app* family triple mutants. These results suggest that APP and Ric8a-regulated heterotrimeric G proteins form a genetic pathway in microglia that normally suppresses microglial immune activation. In support of this interpretation, studies have shown that heterotrimeric G proteins bind to APP and mediate downstream signaling, in an pathway conserved from invertebrates to vertebrates (Fogel et al., 2014; Milosch et al., 2014; Nishimoto et al., 1993; Ramaker et al., 2013). We also show that monomeric Aβ acts as a potent ligand that activates this pathway and suppresses microglial cytokine transcription and secretion (**Fig. 7**). Previous studies have reported pro-inflammatory effects of Aβ oligomers (Bamberger and Landreth, 2001; Bianca et al., 1999; Casal et al., 2004; Combs et al., 2001; Glass et al., 2010; Halle et al., 2008; Ishii et al., 2000; Lee et al., 1993; Lorton et al., 1996; McDonald et al., 1997; Muehlhauser et al., 2001; Reed-Geaghan et al., 2009; Richard et al., 2008; Tan et al., 1999; Van Muiswinkel et al., 1999; Weldon et al., 1998). We show that Aβ monomers are instead anti-inflammatory and suppress microglial activation through an APP and Ric8a-mediated pathway. Thus, our results uncover not only a novel activity of monomeric Aβ in the regulation of brain innate immunity but also the signaling mechanisms underlying this regulation. In parallel to these observations, studies show that Aβ monomers are also protective, while Aβ oligomers are toxic to neurons (Giuffrida et al., 2009; Plant et al., 2003; Ramsden et al., 2002). This suggests that a range of cell types may be similarly regulated by Aβ monomers and oligomers in the nervous system.

APP is an ancient molecule that exists in the earliest multicellular eukaryotes, before the evolution of specialized immune cells (Miller et al., 2007; Tharp and Sarkar, 2013). Several lines of evidence suggest that its first biological function may be linked to early forms of immunity. For example, APP has been found to possess ferroxidase activity that promotes iron export from cells (Duce et al., 2010). Iron export is one of the most ancient mechanisms of nutritional immunity activated by infected cells to deprive intracellular pathogens of essential elements (Soares and Weiss, 2015). Aβ is structurally similar to antimicrobial peptides and possesses strong antimicrobial activity (Kumar et al., 2016), which is another ancient immune mechanism. As specialized immune cells evolve, APP function appears to have diverged in immune and non-immune cells. However, in non-immune cells, there is evidence that APP may have retained its function in these ancient mechanisms of immunity. For example, exposure of several neural lineage cell types including neurons and astrocytes to cytokines has been found to drastically increase their expression and cleavage of APP (Blasko et al., 1999; Buxbaum et al., 1992; Del Bo et al., 1995; He et al., 2007; Hur et al., 2020; Liao et al., 2004; Ohyagi and Tabira, 1993; Rogers et al., 1999). This is consistent with the potential activation of APP/Aβ-mediated innate immunity. We find that Aβ monomers are a potent ligand for suppressing microglial activation. Others have found that Aβ monomers inhibit T cell activation in the periphery (Grant et al., 2012). This suggests that besides its immediate role in innate immune response, Aβ may also act as a negative feedback signal that restrains immune response and as a result potentially facilitate the maintenance of immune homeostasis. Indeed, like Aβ, the prototypic mammalian antimicrobial peptide LL37 has also been found to possesses an immune modulatory activity that down-regulates pro-inflammatory signaling (Mookherjee et al., 2006). Pancortin, another APP-binding ligand (Rice et al., 2012) also belongs to an olfactomedin protein family that similarly acts as a negative feedback signal in innate immunity (Liu et al., 2010). Thus, our results suggest that Aβ monomers may normally play part in the feedback mechanisms that maintain innate immune homeostasis in the brain.

### Aβ-regulated microglial homeostasis and Alzheimer’s disease

The accumulation of Aβ-containing plaques is an early pathological hallmark of Alzheimer’s disease associated with brain microglial activation and followed by formation of tau-containing neurofibrillary tangles that lead to neurodegeneration and cognitive decline (Shi and Holtzman, 2018; Yeh et al., 2017). There is mounting evidence from different fields of research that one of the normal biological functions of Aβ may be to mediate neural activity-dependent axon/synapse competition in the brain and that perturbation of the delicate balance of Aβ monomers and oligomers that mediate this competition may potentially underlie the development of Alzheimer’s disease (Huang, 2023). A relevant point is the opposite ways in which Aβ oligomers and monomers regulate microglial and other glial activity in the brain. Large numbers of studies have shown microglia and other glia play pivotal roles in activity-dependent axon/synapse competition, in which they secrete cytokines, engage in phagocytosis, and promote the elimination of axons/synapses (Li et al., 2020; Madore et al., 2020; Schafer et al., 2012; Vainchtein et al., 2018). Other studies have shown that competing axons/synapses produce Aβ oligomers in an activity-dependent manner, which inhibit the function of competitors and promoting their elimination through receptor dependent and independent mechanisms (Huang, 2023). It is long known that Aβ oligomers strongly activate glial cell types including microglia and astrocytes. Thus, it appears that Aβ oligomers produced by competing axons/synapses mobilize neuronal and glial pathways for competitor elimination in a coordinate manner. Besides competitor elimination, competing axons/synapses must also protect themselves in this process. In cell competition, competing cells activate a multitude of mechanisms to keep Aβ-like agents in their immediate vicinity in the monomer conformation, a conformation frequently associated with strong trophic activity, to protect themselves. In axon/synapse competition, Aβ monomers, which can activate trophic signaling and protect neurons (Giuffrida et al., 2009; Plant et al., 2003; Ramsden et al., 2002), may thus potentially play a similar role. Besides activating neurotrophic signaling, we have shown here that Aβ monomers also potently suppress microglial activation. Thus, like Aβ oligomers, Aβ monomers also appear to engage parallel neuronal and glial mechanisms while performing their protective role. By inhibiting glial activation, which is essential for axon/synapse pruning, Aβ monomers may potentially remove another player in axon/synapse elimination and as such promote protection. Together, these observations suggest that Aβ oligomers and monomers potentially mobilize a coordinated cohort of mechanisms to regulate neuronal as well as glial activity in the processes of brain development and plasticity. Because of the intricate interactions and the wide range of targets, the perturbation of these mechanisms also provides a potential explanation for the pleiotropic effects of Aβ aggregate and plaque formation in Alzheimer’s disease brains (Huang, 2023). For example, studies have shown that chronic neuroinflammation, as may be induced by Aβ aggregation, not only leads to tau hyperphosphorylation and aggregation, but may also compromise neuronal lysosome function that normally facilitate tau clearance (Curnock et al., 2023; Guerrero et al., 2021). Thus, the regulation of microglial and other glial activity, by the balance between Aβ oligomers and monomers, potentially provides a link between amyloidosis and tauopathy in Alzheimer’s disease.

## EXPERIMENTAL PROCEDURES

### Generation of *ric8a* conditional allele

Standard molecular biology techniques were employed for generating the conditional *ric-8a* allele. Briefly, genomic fragments, of 4.5 and 2.5 kilobases and flanking exons 2-4 of the *ric-8a* locus at the 5’ and 3’ side, respectively, were isolated by PCR using high fidelity polymerases. Targeting plasmid was constructed by flanking the genomic fragment containing exons 2-4 with two loxP sites together with a *neomycin* positive selection cassette, followed by 5’ and 3’ genomic fragments as homologous recombination arms and a *pgk-DTA* gene as a negative selection cassette. ES cell clones were screened by Southern blot analysis using external probes at 5’ and 3’ sides. For derivation of conditional allele, the *neomycin* cassette was removed by crossing to an *actin-flpe* transgenic line after blastocyst injection and germ line transmission. The primer set for genotyping *ric-8a* conditional allele, which produces a wildtype band of ∼110bp and a mutant band of ∼200bp, is: 5’-cctagttgtgaatcagaagcacttg-3’ and 5’-gccatacctgagttacctaggc-3’. Animals homozygous for the conditional *ric-8a* allele are viable and fertile, without obvious phenotypes.

### Mouse breeding and pharmacology

*emx1-cre, nestin*-*cre, foxg1-cre, cx3cr1-cre, floxed app* as well as the *BAT-lacZ* reporter mouse lines were purchased from the Jackson Lab. *nex-cre* and *wnt3a-cre* were as published (Goebbels et al., 2006; Yoshida et al., 2006). *cre* transgenes were introduced individually into the *ric8a or app* conditional mutant background for phenotypic analyses and *ric8a or app* homozygotes without *cre* as well as heterozygotes with *cre* (littermates) were both analyzed as controls. For BB94 and MMP9/13 inhibitor injection, pregnant females were treated daily from E12.5 to E14.5 at 30 μg (BB94) or 37.5 μg (MMP9/13 inhibitor) per g of body weight. For dorsomorphin and S3I-201 injection, pregnant females were treated on E12.5 at 7.5 and 25 μg per g of body weight, respectively. For sham treatment, pregnant females were treated on E12.5 with 100 μls of DMSO. BrdU was injected at 100 μg per g of body weight, and embryos were collected 4 hours later for cell proliferation analysis, or alternatively, pups were sacrificed at P5 for neuronal migration analysis and at P17 for other analysis. For LPS treatment, pregnant females were injected intraperitoneally with 400ng (*ric8a* genetic background) or 150ng (*app* genetic background) LPS per g of body weight on both E11.5 and E12.5. Animal use was in accordance with institutional guidelines.

### Immunohistochemistry

Vibratome sections from brains fixed in 4% paraformaldehyde were used. The following primary antibodies were used at respective dilutions/concentrations: mouse anti-BrdU supernatant (clone G3G4, Developmental Studies Hybridoma Bank (DSHB), University of Iowa, IA; 1:40), mouse anti-RC2 supernatant (DSHB; 1:10), mouse anti-Nestin supernatant (DSHB; 1:20), mouse anti-Vimentin supernatant (DSHB; 1:10), mouse anti-Pax6 supernatant (DSHB; 1:20), moue anti-Reelin (Millipore, 1:500), mouse anti-chondroitin sulfate (CS-56, Sigma, 1:100), rat anti-Ctip2 (Abcam, 1:500), rabbit anti-phospho Histone H3 (Ser10) (Millipore; 1:400), rabbit anti-Cux1 (CDP) (Santa Cruz; 1:100), rabbit anti-laminin (Sigma; 1:2000), rabbit anti-GFAP (Dako;1:1000), rabbit anti-ALDH1L1 (Abcam, 1:500), rabbit anti-MMP9 (Abcam, 1:1000), goat anti-MMP2 (R&D Systems; 5 μg/ml), rabbit anti-Calretinin (Chemicon, 1:2000), mouse anti-S100β (Thermo Scientific; 1:100), rabbit anti-S100β (Thermo Scientific; 1:200), and rabbit anti-phospho-Smad1/5 (Ser463/465) (41D10; Cell Signaling, 1:200). FITC and Cy3 conjugated secondary antibodies were purchased from Jackson ImmunoResearch Laboratories (West Grove, PA). Peroxidase conjugated secondary antibodies were purchased from Santa Cruz Biotech. Staining procedures were performed as described previously (Huang et al., 2006), except for anti-Ric-8a, MMP9, and phospho-Smad1/5 staining, in which a tyramide signal amplification (TSA) plus Cy3 kit (PerkinElmer, Waltham, MA) was used per manufacturer’s instruction. Sections were mounted with Fluoromount G medium (Southern Biotech, Birmingham, AB) and analyzed under a Nikon *eclipse* Ti microscope or an Olympus confocal microscope.

### Microglia culture and assay

Cerebral hemispheres were dissected from individual neonates, mechanically dissociated, split into 3-4 wells each and cultured in DMEM-F12 (Lonza) containing 10% fetal bovine serum (FBS) (Invitrogen). Microglial cells were harvested by light trypsinization that removes astroglial sheet on day 13-15. For experiments other than assaying IL-1β secretion, microglia were treated with LPS at 20ng/ml for 3 hours or at 5ng/ml overnight and, if applicable, DMSO or Aβ40 (ApexBio and Genscript) was applied at the same time as LPS. For assaying IL-1β secretion, microglia were primed with LPS at 200ng/ml for 5-6 hours before treatment with 3mM ATP for 15 minutes. In these experiments, DMSO or Aβ40 was applied at the same time as ATP if applicable. Supernatants were collected and used for cytokine ELISA assays per manufacturer’s instructions (Biolegend). Total RNAs were prepared from collected cells using Trizol (Invitrogen) and cDNAs were synthesized using a High-capacity cDNA reverse transcription kit (Applied Biosystems). Quantitative PCR was performed using a GoTaq qPCR master mix per manufacturer’s instructions (Promega). All gene expression levels were normalized against that of GAPDH.

### Quantitative analysis

The sample size was estimated to be 3-9 animals each genotype (every 4^th^ of 50 μm coronal sections, 7-10 sections each animal) for ectopia analysis, 3-5 animals each genotype (3-4 sections each animal) for immunohistochemical analysis, and 4-6 animals each genotype for gel zymography and Western blot analysis, as has been demonstrated by previous publications to be adequate for similar animal studies. Matching sections were used between controls and mutants. NIS-Elements BR 3.0 software (Nikon) was used for quantifying the numbers and sizes of neuronal ectopia, the numbers of laminin positive debris, as well as the numbers of astrocytes. ImageJ software (NIH) was used for quantifying the intensity of immunostainings. Statistics was performed using Student’s *t* test when comparing two conditions, or one-way ANOVA followed by Tukey’s post hoc test when comparing three or more conditions. All data are represented as means ± s.e.m.

## ACKNOWLEDGMENTS

Z. H. thanks Dr. L. F. Reichardt for supporting the initial generation of *ric8a* mutant ES cells, Dr. E. A. Grove (Chicago) for providing the *wnt3a-cre* strain, the late Dr. B. A. Barres (Stanford) for critical input, and Drs. W. L. Murphy and E. Bresnick (UW-Madison) for access to a plate reader and a qPCR machine. We also thank the late Dr. D. Oertel (UW-Madison) for critical reading and editing and Dr. L Puglielli (UW-Madison) for critical reading of a previous version of the manuscript. This work was supported in part by funds from the Departments of Neurology and Neuroscience, UW-Madison, a Basil O’Connor award from the March of Dimes foundation, and an NIH grant NS076729 to Z.H.

## AUTHOR CONTRIBUTIONS

Z. H. designed experiments, generated *ric8a* conditional ES cells, performed microglial and related experiments, and wrote the manuscript. H. J. K., S.M., D. S., and Z. H. performed other experiments and analyzed data.

## REFERENCES

Afshar, K., Willard, F.S., Colombo, K., Johnston, C.A., McCudden, C.R., Siderovski, D.P., and Gonczy, P. (2004). RIC-8 is required for GPR-1/2-dependent Galpha function during asymmetric division of C. elegans embryos. Cell 119, 219–230.

Bamberger, M.E., and Landreth, G.E. (2001). Microglial interaction with beta-amyloid: implications for the pathogenesis of Alzheimer’s disease. Microsc Res Tech 54, 59–70.

Beggs, H.E., Schahin-Reed, D., Zang, K., Goebbels, S., Nave, K.A., Gorski, J., Jones, K.R., Sretavan, D., and Reichardt, L.F. (2003). FAK deficiency in cells contributing to the basal lamina results in cortical abnormalities resembling congenital muscular dystrophies. Neuron 40, 501–514.

Belvindrah, R., Nalbant, P., Ding, S., Wu, C., Bokoch, G.M., and Muller, U. (2006). Integrin-linked kinase regulates Bergmann glial differentiation during cerebellar development. Molecular and cellular neurosciences 33, 109–125.

Bianca, V.D., Dusi, S., Bianchini, E., Dal Pra, I., and Rossi, F. (1999). beta-amyloid activates the O-2 forming NADPH oxidase in microglia, monocytes, and neutrophils. A possible inflammatory mechanism of neuronal damage in Alzheimer’s disease. The Journal of biological chemistry 274, 15493–15499.

Blasko, I., Marx, F., Steiner, E., Hartmann, T., and Grubeck-Loebenstein, B. (1999). TNFalpha plus IFNgamma induce the production of Alzheimer beta-amyloid peptides and decrease the secretion of APPs. FASEB J 13, 63–68.

Buxbaum, J.D., Oishi, M., Chen, H.I., Pinkas-Kramarski, R., Jaffe, E.A., Gandy, S.E., and Greengard, P. (1992). Cholinergic agonists and interleukin 1 regulate processing and secretion of the Alzheimer beta/A4 amyloid protein precursor. Proceedings of the National Academy of Sciences of the United States of America 89, 10075–10078.

Casal, C., Serratosa, J., and Tusell, J.M. (2004). Effects of beta-AP peptides on activation of the transcription factor NF-kappaB and in cell proliferation in glial cell cultures. Neurosci Res 48, 315–323.

Chamberlain, L.M., Holt-Casper, D., Gonzalez-Juarrero, M., and Grainger, D.W. (2015). Extended culture of macrophages from different sources and maturation results in a common M2 phenotype. J Biomed Mater Res A 103, 2864–2874.

Colonna, M., and Butovsky, O. (2017). Microglia Function in the Central Nervous System During Health and Neurodegeneration. Annu Rev Immunol 35, 441–468.

Combs, C.K., Karlo, J.C., Kao, S.C., and Landreth, G.E. (2001). beta-Amyloid stimulation of microglia and monocytes results in TNFalpha-dependent expression of inducible nitric oxide synthase and neuronal apoptosis. The Journal of neuroscience: the official journal of the Society for Neuroscience 21, 1179–1188.

Couwenbergs, C., Spilker, A.C., and Gotta, M. (2004). Control of embryonic spindle positioning and Galpha activity by C. elegans RIC-8. Current biology: CB 14, 1871–1876.

Covarrubias, A.J., Aksoylar, H.I., Yu, J., Snyder, N.W., Worth, A.J., Iyer, S.S., Wang, J., Ben-Sahra, I., Byles, V., Polynne-Stapornkul, T., et al. (2016). Akt-mTORC1 signaling regulates Acly to integrate metabolic input to control of macrophage activation. eLife 5.

Curnock, R., Yalci, K., Palmfeldt, J., Jaattela, M., Liu, B., and Carroll, B. (2023). TFEB-dependent lysosome biogenesis is required for senescence. The EMBO journal 42, e111241.

David, N.B., Martin, C.A., Segalen, M., Rosenfeld, F., Schweisguth, F., and Bellaiche, Y. (2005). Drosophila Ric-8 regulates Galphai cortical localization to promote Galphai-dependent planar orientation of the mitotic spindle during asymmetric cell division. Nature cell biology 7, 1083–1090.

De Strooper, B., Iwatsubo, T., and Wolfe, M.S. (2012). Presenilins and gamma-secretase: structure, function, and role in Alzheimer Disease. Cold Spring Harb Perspect Med 2, a006304.

Del Bo, R., Angeretti, N., Lucca, E., De Simoni, M.G., and Forloni, G. (1995). Reciprocal control of inflammatory cytokines, IL-1 and IL-6, and beta-amyloid production in cultures. Neuroscience letters 188, 70–74.

Duce, J.A., Tsatsanis, A., Cater, M.A., James, S.A., Robb, E., Wikhe, K., Leong, S.L., Perez, K., Johanssen, T., Greenough, M.A., et al. (2010). Iron-export ferroxidase activity of beta-amyloid precursor protein is inhibited by zinc in Alzheimer’s disease. Cell 142, 857–867.

Fogel, H., Frere, S., Segev, O., Bharill, S., Shapira, I., Gazit, N., O’Malley, T., Slomowitz, E., Berdichevsky, Y., Walsh, D.M., et al. (2014). APP homodimers transduce an amyloid-beta-mediated increase in release probability at excitatory synapses. Cell reports 7, 1560–1576.

Franco, S.J., and Muller, U. (2013). Shaping our minds: stem and progenitor cell diversity in the mammalian neocortex. Neuron 77, 19–34.

Gabay, M., Pinter, M.E., Wright, F.A., Chan, P., Murphy, A.J., Valenzuela, D.M., Yancopoulos, G.D., and Tall, G.G. (2011). Ric-8 proteins are molecular chaperones that direct nascent G protein alpha subunit membrane association. Science signaling 4, ra79.

Ginhoux, F., Greter, M., Leboeuf, M., Nandi, S., See, P., Gokhan, S., Mehler, M.F., Conway, S.J., Ng, L.G., Stanley, E.R., et al. (2010). Fate mapping analysis reveals that adult microglia derive from primitive macrophages. Science 330, 841–845.

Giuffrida, M.L., Caraci, F., Pignataro, B., Cataldo, S., De Bona, P., Bruno, V., Molinaro, G., Pappalardo, G., Messina, A., Palmigiano, A., et al. (2009). Beta-amyloid monomers are neuroprotective. The Journal of neuroscience: the official journal of the Society for Neuroscience 29, 10582–10587.

Glass, C.K., Saijo, K., Winner, B., Marchetto, M.C., and Gage, F.H. (2010). Mechanisms underlying inflammation in neurodegeneration. Cell 140, 918–934.

Goebbels, S., Bormuth, I., Bode, U., Hermanson, O., Schwab, M.H., and Nave, K.A. (2006). Genetic targeting of principal neurons in neocortex and hippocampus of NEX-Cre mice. Genesis 44, 611–621.

Gorski, J.A., Talley, T., Qiu, M., Puelles, L., Rubenstein, J.L., and Jones, K.R. (2002). Cortical excitatory neurons and glia, but not GABAergic neurons, are produced in the Emx1-expressing lineage. The Journal of neuroscience: the official journal of the Society for Neuroscience 22, 6309–6314.

Grant, J.L., Ghosn, E.E., Axtell, R.C., Herges, K., Kuipers, H.F., Woodling, N.S., Andreasson, K., Herzenberg, L.A., Herzenberg, L.A., and Steinman, L. (2012). Reversal of paralysis and reduced inflammation from peripheral administration of beta-amyloid in TH1 and TH17 versions of experimental autoimmune encephalomyelitis. Sci Transl Med 4, 145ra105.

Graus-Porta, D., Blaess, S., Senften, M., Littlewood-Evans, A., Damsky, C., Huang, Z., Orban, P., Klein, R., Schittny, J.C., and Muller, U. (2001). Beta1-class integrins regulate the development of laminae and folia in the cerebral and cerebellar cortex. Neuron 31, 367–379.

Guenette, S., Chang, Y., Hiesberger, T., Richardson, J.A., Eckman, C.B., Eckman, E.A., Hammer, R.E., and Herz, J. (2006). Essential roles for the FE65 amyloid precursor protein-interacting proteins in brain development. The EMBO journal 25, 420–431.

Guerrero, A., De Strooper, B., and Arancibia-Carcamo, I.L. (2021). Cellular senescence at the crossroads of inflammation and Alzheimer’s disease. Trends Neurosci 44, 714–727.

Haass, C., Kaether, C., Thinakaran, G., and Sisodia, S. (2012). Trafficking and proteolytic processing of APP. Cold Spring Harb Perspect Med 2, a006270.

Halle, A., Hornung, V., Petzold, G.C., Stewart, C.R., Monks, B.G., Reinheckel, T., Fitzgerald, K.A., Latz, E., Moore, K.J., and Golenbock, D.T. (2008). The NALP3 inflammasome is involved in the innate immune response to amyloid-beta. Nat Immunol 9, 857–865.

Hampoelz, B., Hoeller, O., Bowman, S.K., Dunican, D., and Knoblich, J.A. (2005). Drosophila Ric-8 is essential for plasma-membrane localization of heterotrimeric G proteins. Nat Cell Biol 7, 1099–1105.

Hanania, R., Sun, H.S., Xu, K., Pustylnik, S., Jeganathan, S., and Harrison, R.E. (2012). Classically activated macrophages use stable microtubules for matrix metalloproteinase-9 (MMP-9) secretion. The Journal of biological chemistry 287, 8468–8483.

Hansen, D.V., Rubenstein, J.L., and Kriegstein, A.R. (2011). Deriving excitatory neurons of the neocortex from pluripotent stem cells. Neuron 70, 645–660.

Hattori, Y., Kato, D., Murayama, F., Koike, S., Asai, H., Yamasaki, A., Naito, Y., Kawaguchi, A., Konishi, H., Prinz, M., et al. (2023). CD206(+) macrophages transventricularly infiltrate the early embryonic cerebral wall to differentiate into microglia. Cell reports 42, 112092.

He, P., Zhong, Z., Lindholm, K., Berning, L., Lee, W., Lemere, C., Staufenbiel, M., Li, R., and Shen, Y. (2007). Deletion of tumor necrosis factor death receptor inhibits amyloid beta generation and prevents learning and memory deficits in Alzheimer’s mice. J Cell Biol 178, 829–841.

Hebert, J.M., and McConnell, S.K. (2000). Targeting of cre to the Foxg1 (BF-1) locus mediates loxP recombination in the telencephalon and other developing head structures. Developmental biology 222, 296–306.

Herms, J., Anliker, B., Heber, S., Ring, S., Fuhrmann, M., Kretzschmar, H., Sisodia, S., and Muller, U. (2004). Cortical dysplasia resembling human type 2 lissencephaly in mice lacking all three APP family members. The EMBO journal 23, 4106–4115.

Huang, Z. (2023). A Function of Amyloid-beta in Mediating Activity-Dependent Axon/Synapse Competition May Unify Its Roles in Brain Physiology and Pathology. J Alzheimers Dis 92, 29–57.

Huang, Z., Shimazu, K., Woo, N.H., Zang, K., Muller, U., Lu, B., and Reichardt, L.F. (2006). Distinct roles of the beta 1-class integrins at the developing and the mature hippocampal excitatory synapse. J Neurosci 26, 11208–11219.

Hur, J.Y., Frost, G.R., Wu, X., Crump, C., Pan, S.J., Wong, E., Barros, M., Li, T., Nie, P., Zhai, Y., et al. (2020). The innate immunity protein IFITM3 modulates gamma-secretase in Alzheimer’s disease. Nature 586, 735–740.

Ishii, K., Muelhauser, F., Liebl, U., Picard, M., Kuhl, S., Penke, B., Bayer, T., Wiessler, M., Hennerici, M., Beyreuther, K., et al. (2000). Subacute NO generation induced by Alzheimer’s beta-amyloid in the living brain: reversal by inhibition of the inducible NO synthase. FASEB J 14, 1485–1489.

Iwasawa, E., Brown, F.N., Shula, C., Kahn, F., Lee, S.H., Berta, T., Ladle, D.R., Campbell, K., Mangano, F.T., and Goto, J. (2022). The Anti-Inflammatory Agent Bindarit Attenuates the Impairment of Neural Development through Suppression of Microglial Activation in a Neonatal Hydrocephalus Mouse Model. The Journal of neuroscience: the official journal of the Society for Neuroscience 42, 1820–1844.

Jeong, S.J., Luo, R., Singer, K., Giera, S., Kreidberg, J., Kiyozumi, D., Shimono, C., Sekiguchi, K., and Piao, X. (2013). GPR56 functions together with alpha3beta1 integrin in regulating cerebral cortical development. PLoS One 8, e68781.

Kumar, D.K., Choi, S.H., Washicosky, K.J., Eimer, W.A., Tucker, S., Ghofrani, J., Lefkowitz, A., McColl, G., Goldstein, L.E., Tanzi, R.E., et al. (2016). Amyloid-beta peptide protects against microbial infection in mouse and worm models of Alzheimer’s disease. Sci Transl Med 8, 340ra372.

Lee, S.C., Liu, W., Dickson, D.W., Brosnan, C.F., and Berman, J.W. (1993). Cytokine production by human fetal microglia and astrocytes. Differential induction by lipopolysaccharide and IL-1 beta. Journal of immunology 150, 2659–2667.

LeVine, H., 3rd (2004). Alzheimer’s beta-peptide oligomer formation at physiologic concentrations. Anal Biochem 335, 81–90.

Li, Q., and Barres, B.A. (2018). Microglia and macrophages in brain homeostasis and disease. Nat Rev Immunol 18, 225–242.

Li, S., Jin, Z., Koirala, S., Bu, L., Xu, L., Hynes, R.O., Walsh, C.A., Corfas, G., and Piao, X. (2008). GPR56 regulates pial basement membrane integrity and cortical lamination. The Journal of neuroscience: the official journal of the Society for Neuroscience 28, 5817–5826.

Li, T., Chiou, B., Gilman, C.K., Luo, R., Koshi, T., Yu, D., Oak, H.C., Giera, S., Johnson-Venkatesh, E., Muthukumar, A.K., et al. (2020). A splicing isoform of GPR56 mediates microglial synaptic refinement via phosphatidylserine binding. The EMBO journal 39, e104136.

Liao, Y.F., Wang, B.J., Cheng, H.T., Kuo, L.H., and Wolfe, M.S. (2004). Tumor necrosis factor-alpha, interleukin-1beta, and interferon-gamma stimulate gamma-secretase-mediated cleavage of amyloid precursor protein through a JNK-dependent MAPK pathway. The Journal of biological chemistry 279, 49523–49532.

Liu, W., Yan, M., Liu, Y., Wang, R., Li, C., Deng, C., Singh, A., Coleman, W.G., Jr., and Rodgers, G.P. (2010). Olfactomedin 4 down-regulates innate immunity against Helicobacter pylori infection. Proceedings of the National Academy of Sciences of the United States of America 107, 11056–11061.

Lorton, D., Kocsis, J.M., King, L., Madden, K., and Brunden, K.R. (1996). beta-Amyloid induces increased release of interleukin-1 beta from lipopolysaccharide-activated human monocytes. Journal of neuroimmunology 67, 21–29.

Ma, S., Kwon, H.J., and Huang, Z. (2012). Ric-8a, a guanine nucleotide exchange factor for heterotrimeric G proteins, regulates bergmann glia-basement membrane adhesion during cerebellar foliation. The Journal of neuroscience: the official journal of the Society for Neuroscience 32, 14979–14993.

Ma, S., Santhosh, D., Kumar, T.P., and Huang, Z. (2017). A Brain-Region-Specific Neural Pathway Regulating Germinal Matrix Angiogenesis. Developmental cell 41, 366–381 e364.

Madore, C., Leyrolle, Q., Morel, L., Rossitto, M., Greenhalgh, A.D., Delpech, J.C., Martinat, M., Bosch-Bouju, C., Bourel, J., Rani, B., et al. (2020). Essential omega-3 fatty acids tune microglial phagocytosis of synaptic elements in the mouse developing brain. Nat Commun 11, 6133.

Marin, O., and Rubenstein, J.L. (2003). Cell migration in the forebrain. Annual review of neuroscience 26, 441–483.

McDonald, D.R., Brunden, K.R., and Landreth, G.E. (1997). Amyloid fibrils activate tyrosine kinase-dependent signaling and superoxide production in microglia. The Journal of neuroscience: the official journal of the Society for Neuroscience 17, 2284–2294.

Miller, D.J., Hemmrich, G., Ball, E.E., Hayward, D.C., Khalturin, K., Funayama, N., Agata, K., and Bosch, T.C. (2007). The innate immune repertoire in cnidaria--ancestral complexity and stochastic gene loss. Genome Biol 8, R59.

Milosch, N., Tanriover, G., Kundu, A., Rami, A., Francois, J.C., Baumkotter, F., Weyer, S.W., Samanta, A., Jaschke, A., Brod, F., et al. (2014). Holo-APP and G-protein-mediated signaling are required for sAPPalpha-induced activation of the Akt survival pathway. Cell death & disease 5, e1391.

Mookherjee, N., Brown, K.L., Bowdish, D.M., Doria, S., Falsafi, R., Hokamp, K., Roche, F.M., Mu, R., Doho, G.H., Pistolic, J., et al. (2006). Modulation of the TLR-mediated inflammatory response by the endogenous human host defense peptide LL-37. Journal of immunology 176, 2455–2464.

Moore, S.A., Saito, F., Chen, J., Michele, D.E., Henry, M.D., Messing, A., Cohn, R.D., Ross-Barta, S.E., Westra, S., Williamson, R.A., et al. (2002). Deletion of brain dystroglycan recapitulates aspects of congenital muscular dystrophy. Nature 418, 422–425.

Muehlhauser, F., Liebl, U., Kuehl, S., Walter, S., Bertsch, T., and Fassbender, K. (2001). Aggregation-Dependent interaction of the Alzheimer’s beta-amyloid and microglia. Clin Chem Lab Med 39, 313–316.

Muller, U.C., and Zheng, H. (2012). Physiological functions of APP family proteins. Cold Spring Harb Perspect Med 2, a006288.

Niewmierzycka, A., Mills, J., St-Arnaud, R., Dedhar, S., and Reichardt, L.F. (2005). Integrin-linked kinase deletion from mouse cortex results in cortical lamination defects resembling cobblestone lissencephaly. The Journal of neuroscience: the official journal of the Society for Neuroscience 25, 7022–7031.

Nishimoto, I., Okamoto, T., Matsuura, Y., Takahashi, S., Okamoto, T., Murayama, Y., and Ogata, E. (1993). Alzheimer amyloid protein precursor complexes with brain GTP-binding protein G(o). Nature 362, 75–79.

Ohyagi, Y., and Tabira, T. (1993). Effect of growth factors and cytokines on expression of amyloid beta protein precursor mRNAs in cultured neural cells. Brain research Molecular brain research 18, 127–132.

Plant, L.D., Boyle, J.P., Smith, I.F., Peers, C., and Pearson, H.A. (2003). The production of amyloid beta peptide is a critical requirement for the viability of central neurons. The Journal of neuroscience: the official journal of the Society for Neuroscience 23, 5531–5535.

Qin, H., Yeh, W.I., De Sarno, P., Holdbrooks, A.T., Liu, Y., Muldowney, M.T., Reynolds, S.L., Yanagisawa, L.L., Fox, T.H., 3rd, Park, K., et al. (2012). Signal transducer and activator of transcription-3/suppressor of cytokine signaling-3 (STAT3/SOCS3) axis in myeloid cells regulates neuroinflammation. Proceedings of the National Academy of Sciences of the United States of America 109, 5004–5009.

Ramaker, J.M., Swanson, T.L., and Copenhaver, P.F. (2013). Amyloid precursor proteins interact with the heterotrimeric G protein Go in the control of neuronal migration. The Journal of neuroscience: the official journal of the Society for Neuroscience 33, 10165–10181.

Ramsden, M., Henderson, Z., and Pearson, H.A. (2002). Modulation of Ca2+ channel currents in primary cultures of rat cortical neurones by amyloid beta protein (1-40) is dependent on solubility status. Brain research 956, 254–261.

Reed-Geaghan, E.G., Savage, J.C., Hise, A.G., and Landreth, G.E. (2009). CD14 and toll-like receptors 2 and 4 are required for fibrillar A{beta}-stimulated microglial activation. The Journal of neuroscience: the official journal of the Society for Neuroscience 29, 11982–11992.

Rice, H.C., Townsend, M., Bai, J., Suth, S., Cavanaugh, W., Selkoe, D.J., and Young-Pearse, T.L. (2012). Pancortins interact with amyloid precursor protein and modulate cortical cell migration. Development 139, 3986–3996.

Richard, K.L., Filali, M., Prefontaine, P., and Rivest, S. (2008). Toll-like receptor 2 acts as a natural innate immune receptor to clear amyloid beta 1-42 and delay the cognitive decline in a mouse model of Alzheimer’s disease. The Journal of neuroscience: the official journal of the Society for Neuroscience 28, 5784–5793.

Rogers, J.T., Leiter, L.M., McPhee, J., Cahill, C.M., Zhan, S.S., Potter, H., and Nilsson, L.N. (1999). Translation of the alzheimer amyloid precursor protein mRNA is up-regulated by interleukin-1 through 5’-untranslated region sequences. The Journal of biological chemistry 274, 6421–6431.

Satz, J.S., Ostendorf, A.P., Hou, S., Turner, A., Kusano, H., Lee, J.C., Turk, R., Nguyen, H., Ross-Barta, S.E., Westra, S., et al. (2010). Distinct functions of glial and neuronal dystroglycan in the developing and adult mouse brain. J Neurosci 30, 14560–14572.

Sayed, F.A., Telpoukhovskaia, M., Kodama, L., Li, Y., Zhou, Y., Le, D., Hauduc, A., Ludwig, C., Gao, F., Clelland, C., et al. (2018). Differential effects of partial and complete loss of TREM2 on microglial injury response and tauopathy. Proceedings of the National Academy of Sciences of the United States of America 115, 10172–10177.

Schafer, D.P., Lehrman, E.K., Kautzman, A.G., Koyama, R., Mardinly, A.R., Yamasaki, R., Ransohoff, R.M., Greenberg, M.E., Barres, B.A., and Stevens, B. (2012). Microglia sculpt postnatal neural circuits in an activity and complement-dependent manner. Neuron 74, 691–705.

Selkoe, D.J., and Hardy, J. (2016). The amyloid hypothesis of Alzheimer’s disease at 25 years. EMBO molecular medicine 8, 595–608.

Shaked, G.M., Kummer, M.P., Lu, D.C., Galvan, V., Bredesen, D.E., and Koo, E.H. (2006). Abeta induces cell death by direct interaction with its cognate extracellular domain on APP (APP 597-624). FASEB J 20, 1254–1256.

Shi, Y., and Holtzman, D.M. (2018). Interplay between innate immunity and Alzheimer disease: APOE and TREM2 in the spotlight. Nat Rev Immunol 18, 759–772.

Soares, M.P., and Weiss, G. (2015). The Iron age of host-microbe interactions. EMBO Rep 16, 1482–1500.

Stine, W.B., Jungbauer, L., Yu, C., and LaDu, M.J. (2011). Preparing synthetic Abeta in different aggregation states. Methods Mol Biol 670, 13–32.

Tall, G.G., Krumins, A.M., and Gilman, A.G. (2003). Mammalian Ric-8A (synembryn) is a heterotrimeric Galpha protein guanine nucleotide exchange factor. The Journal of biological chemistry 278, 8356–8362.

Tan, J., Town, T., Paris, D., Mori, T., Suo, Z., Crawford, F., Mattson, M.P., Flavell, R.A., and Mullan, M. (1999). Microglial activation resulting from CD40-CD40L interaction after beta-amyloid stimulation. Science 286, 2352–2355.

Tanzi, R.E. (2012). The genetics of Alzheimer disease. Cold Spring Harb Perspect Med 2.

Tharp, W.G., and Sarkar, I.N. (2013). Origins of amyloid-beta. BMC Genomics 14, 290.

Vainchtein, I.D., Chin, G., Cho, F.S., Kelley, K.W., Miller, J.G., Chien, E.C., Liddelow, S.A., Nguyen, P.T., Nakao-Inoue, H., Dorman, L.C., et al. (2018). Astrocyte-derived interleukin-33 promotes microglial synapse engulfment and neural circuit development. Science 359, 1269–1273.

Van Muiswinkel, F.L., Raupp, S.F., de Vos, N.M., Smits, H.A., Verhoef, J., Eikelenboom, P., and Nottet, H.S. (1999). The amino-terminus of the amyloid-beta protein is critical for the cellular binding and consequent activation of the respiratory burst of human macrophages. Journal of neuroimmunology 96, 121–130.

Waite, A., Brown, S.C., and Blake, D.J. (2012). The dystrophin-glycoprotein complex in brain development and disease. Trends Neurosci 35, 487–496.

Wang, H., Ng, K.H., Qian, H., Siderovski, D.P., Chia, W., and Yu, F. (2005). Ric-8 controls Drosophila neural progenitor asymmetric division by regulating heterotrimeric G proteins. Nat Cell Biol 7, 1091–1098.

Wang, Z., Wang, B., Yang, L., Guo, Q., Aithmitti, N., Songyang, Z., and Zheng, H. (2009). Presynaptic and postsynaptic interaction of the amyloid precursor protein promotes peripheral and central synaptogenesis. The Journal of neuroscience: the official journal of the Society for Neuroscience 29, 10788–10801.

Weldon, D.T., Rogers, S.D., Ghilardi, J.R., Finke, M.P., Cleary, J.P., O’Hare, E., Esler, W.P., Maggio, J.E., and Mantyh, P.W. (1998). Fibrillar beta-amyloid induces microglial phagocytosis, expression of inducible nitric oxide synthase, and loss of a select population of neurons in the rat CNS in vivo. The Journal of neuroscience: the official journal of the Society for Neuroscience 18, 2161–2173.

Yeh, F.L., Hansen, D.V., and Sheng, M. (2017). TREM2, Microglia, and Neurodegenerative Diseases. Trends Mol Med 23, 512–533.

Yona, S., Kim, K.W., Wolf, Y., Mildner, A., Varol, D., Breker, M., Strauss-Ayali, D., Viukov, S., Guilliams, M., Misharin, A., et al. (2013). Fate mapping reveals origins and dynamics of monocytes and tissue macrophages under homeostasis. Immunity 38, 79–91.

Yoshida, M., Assimacopoulos, S., Jones, K.R., and Grove, E.A. (2006). Massive loss of Cajal-Retzius cells does not disrupt neocortical layer order. Development 133, 537–545.

Young-Pearse, T.L., Bai, J., Chang, R., Zheng, J.B., LoTurco, J.J., and Selkoe, D.J. (2007). A critical function for beta-amyloid precursor protein in neuronal migration revealed by in utero RNA interference. The Journal of neuroscience: the official journal of the Society for Neuroscience 27, 14459–14469.

Zhang, Y., Chen, K., Sloan, S.A., Bennett, M.L., Scholze, A.R., O’Keeffe, S., Phatnani, H.P., Guarnieri, P., Caneda, C., Ruderisch, N., et al. (2014). An RNA-sequencing transcriptome and splicing database of glia, neurons, and vascular cells of the cerebral cortex. The Journal of neuroscience: the official journal of the Society for Neuroscience 34, 11929–11947.

