## Supplemental figures 1-7 for "Monomeric amyloid-β inhibits microglial inflammatory activity in the brain via an APP/heterotrimeric G protein-mediated pathway"

### SUPPLEMENTAL INFORMATION

#### Supplementary Figures and Legends

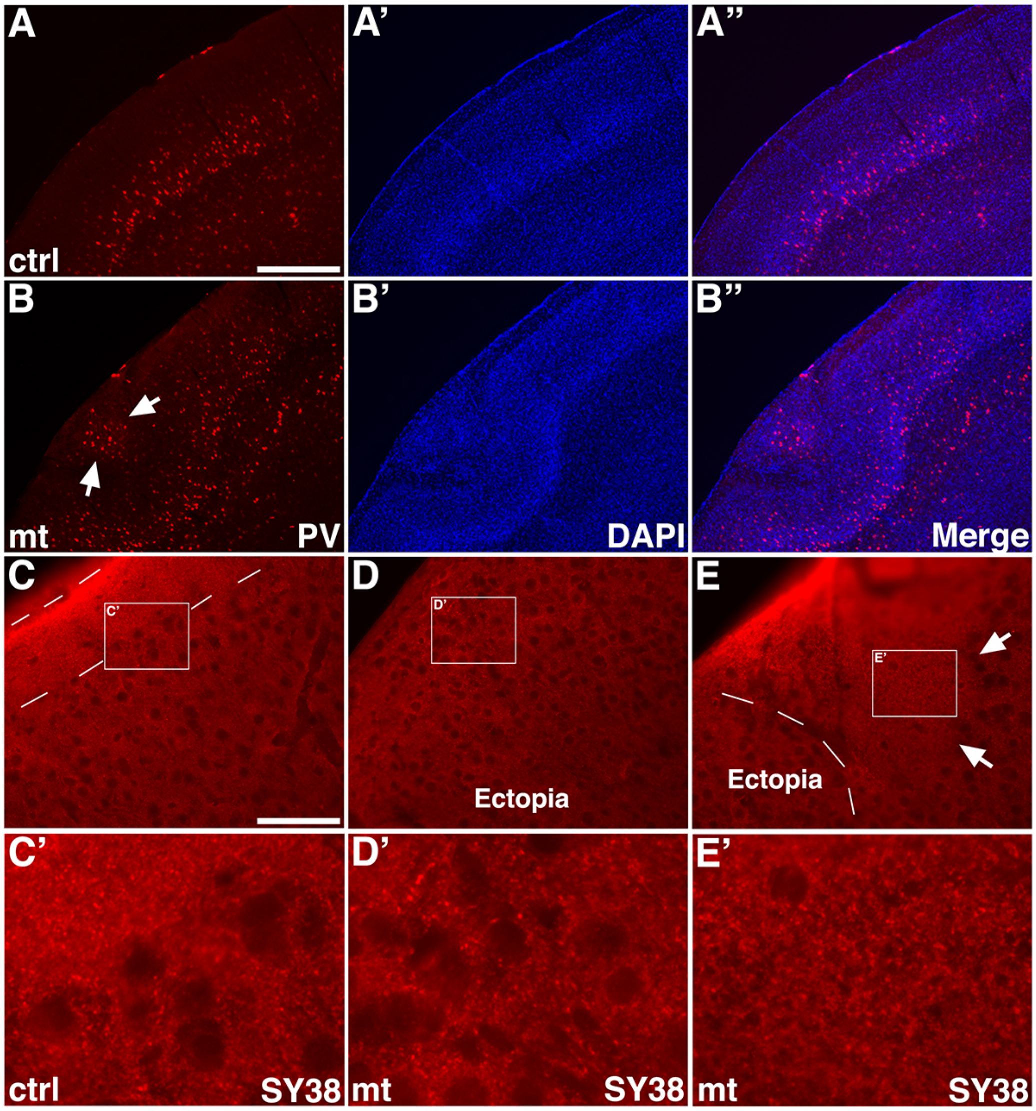


**Figure S1.** ***ric8a/emx1-cre*** **mutation results in disorganization of cortical neural circuitry.** Related to Figure 1.

**(A-B")** Parvalbumin (PV, in red) and nuclear (DAPI, in blue) staining of control (**A-A"**) and mutant (**B-B"**) cortices at P17. Note ectopic PV-positive interneurons in the mutant (arrows in **B**).

(**C-E’**) Synaptic marker SY38 (in red) staining of control (**C** & **C’**) and mutant (**D-E’**) cortices at P17. Outlined areas in (**C-E**) are shown in detail in (**C’, D’, E’**). Region shown in (**D**) is within an ectopia while that in (**E**) is at the border of an ectopia.

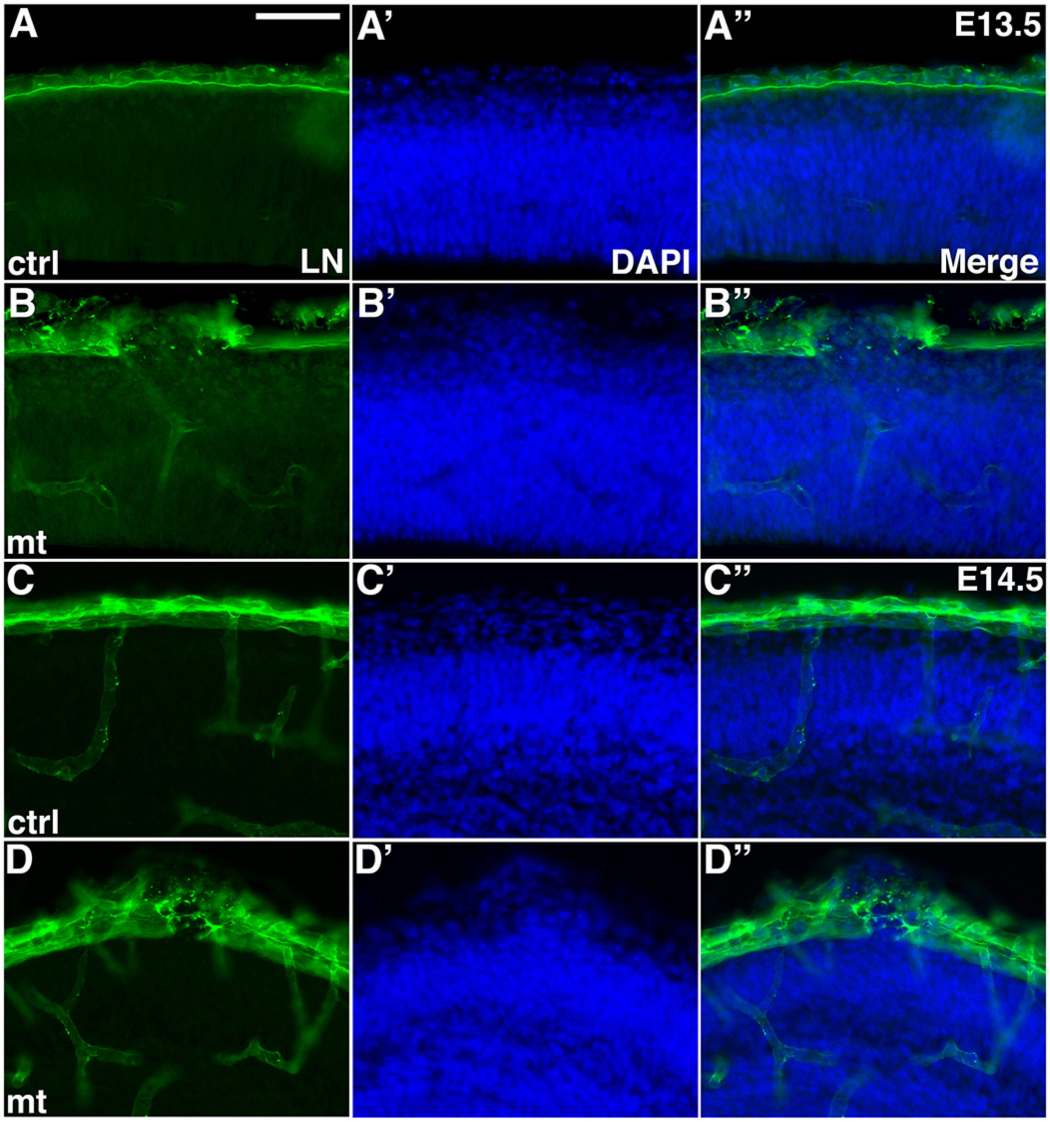


**Figure S2. Neuronal ectopia and basement membrane breach in *ric8a*/*emx1-cre* mutants at E13.5 and E14.5.** Related to Figure 2.

**(A-A”)** Laminin (LN, in green) and nuclear (DAPI, in blue) staining of control cortices at E13.5. A continuous basement membrane is observed at the pia, beneath which cells are well organized in the cortical wall.

**(B-B”)** Laminin and nuclear staining of *ric8a*/*emx1-cre* mutant cortices at E13.5. Although at E13.5 we observe basement membrane defects in the absence of neuronal ectopia (see **Fig. 3A-B”**), neuronal ectopia at this stage is consistently associated with basement membrane breakage.

(**C**-**C”**) Laminin and nuclear staining of control cortices at E14.5.

(**D**-**D”**) Laminin and nuclear staining of *ric-8a/emx1-cre* mutant cortices at E14.5. Neuronal ectopia at E14.5 is also consistently associated with basement membrane breakage.

Scale bar in (**A**), 100 μm for all panels.

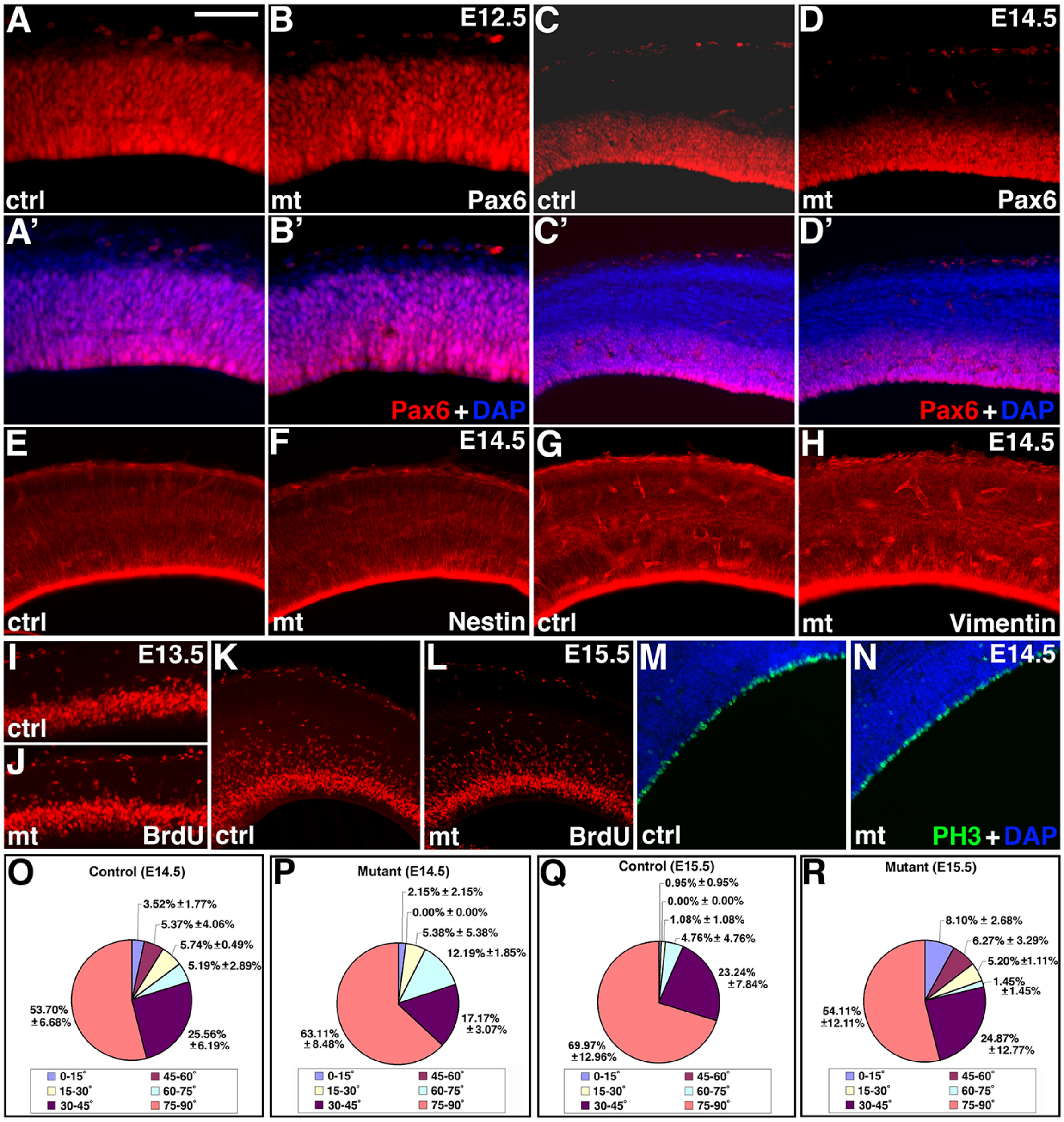


**Figure S3. Cortical radial glial identity and proliferation are overall unaffected in *ric8a/emx1-cre* mutants**. Related to Figure 3.

(**A**-**B'**) Pax6 (in red) and nuclear (DAPI, in blue) staining of control (**A** & **A'**) and mutant (**B** & **B'**) cortices at E12.5.

(**C**-**D’**) Pax6 (in red) and nuclear (DAPI, in blue) staining of control (**C** & **C’**) and mutant (**D** & **D’**) cortices at E14.5. No ectopic Pax6 positive cells were observed at either E12.5 or E14.5.

**(E** & **F**) Nestin (in red) and nuclear (DAPI, in blue) staining of control (**E**) and mutant (**F**) cortices at E14.5.

(**G** & **H**) Vimentin (in red) and nuclear (DAPI, in blue) staining of control (**G**) and mutant (**H**) cortices at E14.5.

**(I** & **J**) BrdU staining (in red) in control (**I**) and mutant (**J**) cortices at E13.5.

(**K** & **L**) BrdU staining (in red) in control (**K**) and mutant (**L**) cortices at E15.5.

(**M** & **N**) Phospho-histone 3 (PH3, in green) and nuclear (DAPI, in blue) staining of control (**M**) and mutant (**N**) cortices at E14.5.

**(O** & **P**) Cleavage plane distribution of radial glial mitosis in control (**O**) and mutant (**P**) cortices at E12.5. No significant differences were observed ($P > 0.4$, n = 3 animals each genotype; 73 cells for controls and 76 cells for mutants).

(**R** & **S**) Cleavage plane distribution of radial glial mitosis in control (**R**) and mutant (**S**) cortices at E14.5. No significant differences were observed ($P > 0.1$, n = 3 animals each genotype; 70 cells for controls and 59 cells for mutants).

Scale bar in (**A**), 100 μm for (**A**-**B’**) and 200 μm for (**C**-**N**).

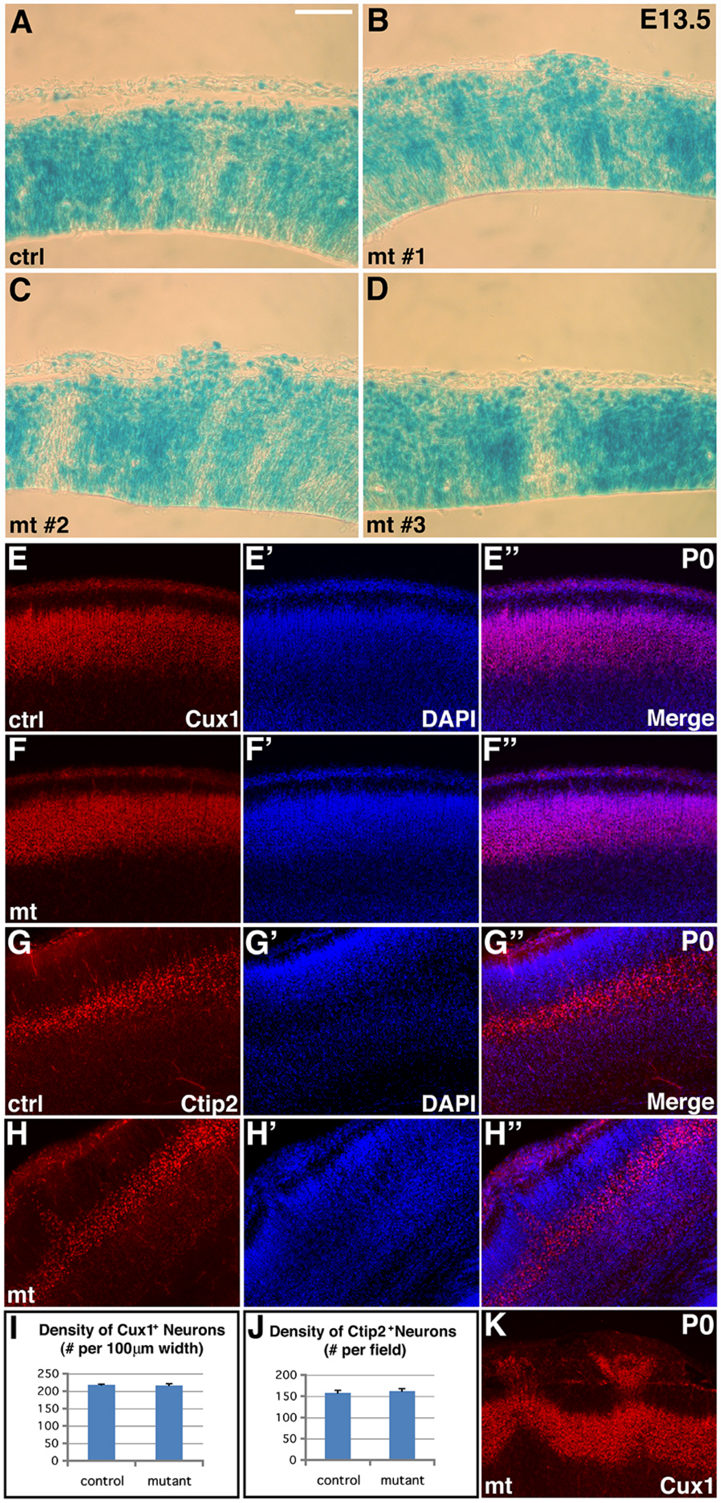


**Figure S4. Wnt pathway activity and lamina-specific neuronal markers are normal in *ric8a*/*emx1-cre* mutant cortices.** Related to Figure 4.

(**A**-**D**) X-gal staining of BAT-lacZ expression in *ric8a*/*emx1-cre* control (**A**) and mutant (**B**-**D**) cortices at E13.5. No obvious differences are observed between controls and three different mutants at this stage.

(**E**-**F**") Cux1 (in red) and nuclear (DAPI, in blue) staining of control (**E-E"**) and mutant (**F-F"**) cortices at P0 in a region without ectopia. No obvious changes in the expression pattern of Cux1, an upper layer neuronal marker, were observed in the mutant cortex, except in areas with ectopia (see panel (**K**)).

(**G**-**H**") Ctip2 (in red) and nuclear (DAPI, in blue) staining of control (**G**-**G**") and mutant (**H**-**H**") cortices at P0. No obvious changes in the expression pattern of Ctip2, a deep layer neuronal marker, were observed in the mutant cortex, except in areas with ectopia.

(**I** & **J**) Quantification of cortical neurons positive for Cux1 (**I**) and Ctip2 (**J**) in matching cortical regions at P0. No significant differences were observed in the density of Cux1 (control, 218.1 ± 1.7 per 100 μm cortical width; mutant, 216.4 ± 4.3 per 100 μm cortical width; $P = 0.36$, n = 12) or Ctip2 (control, 157.8 ± 5.0 per field; mutant, 161.9 ± 5.9 per field; $P = 0.31$, n = 12) positive neurons between controls and mutants.

(K) Cux1 (in red) and nuclear (DAPI, in blue) staining of mutant cortices at P0 in a region with ectopia.

Scale bar in (**A**), 135 μm for (**A-D**) and 200 μm for (**E-H"**).

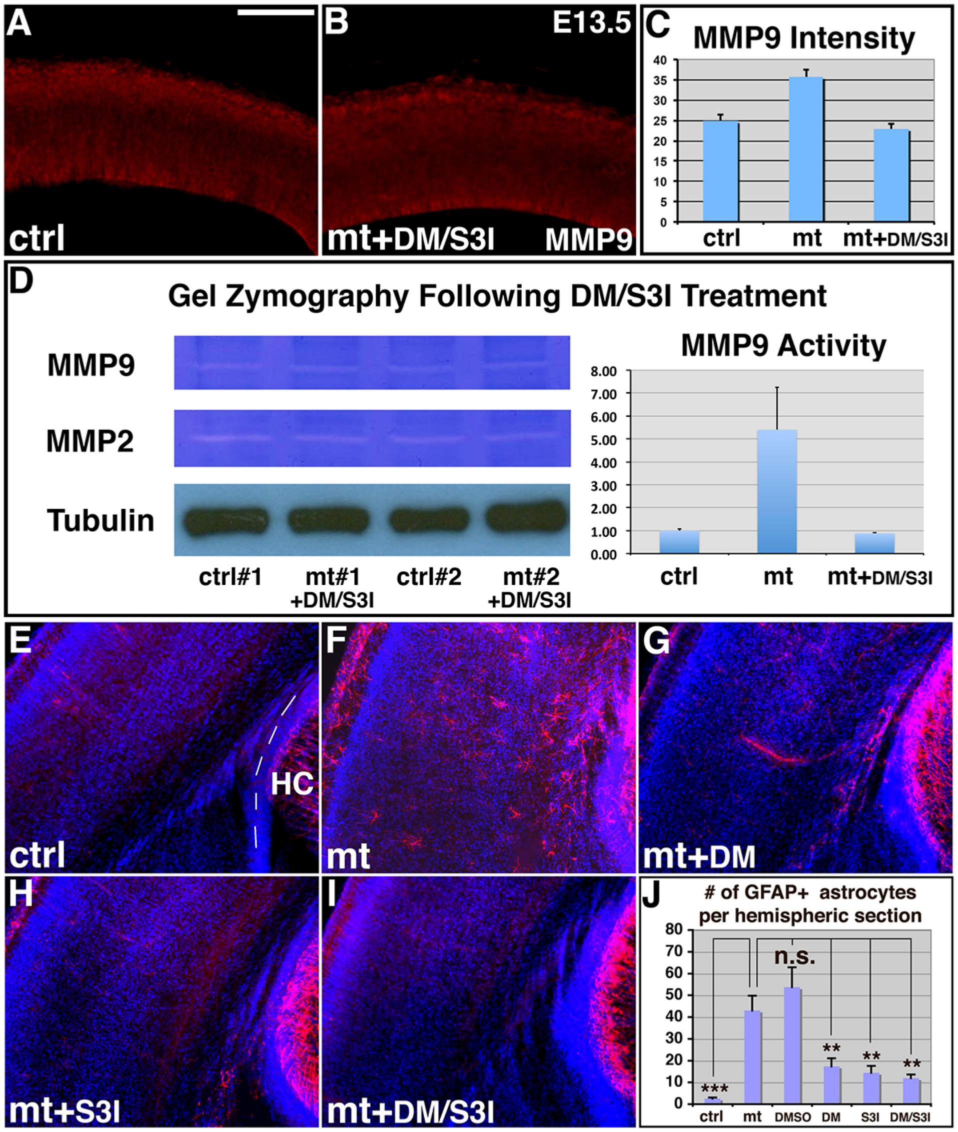


**Figure S5. Suppression of MMP9 expression and gliosis in *ric8a/emx1-cre* mutant cortices by anti-inflammatory inhibitors, dorsomorphin (DM) and S3I-201 (S3I).** Related to Figure 7.

(**A**) MMP9 (in red) staining in control cortices at E13.5.

(**B**) MMP9 (in red) staining in *ric8a/emx1-cre* mutant cortices at E13.5 after DM and S3I dual treatment at E12.5.

(**C**) Quantitative analysis of MMP9 expression. No significant differences are observed in mutants after inhibitor treatment in comparison to controls ($P$ = 0.44, n = 6).

(**D**) Gel zymography of E13.5 control and mutant cortical lysates following DM and S3I treatment at E12.5. Similar levels of MMP9 are observed between controls and mutants. Quantification also showed no significant differences in normalized MMP9 levels ($P$ = 0.46, n = 4).

**(E-I**) GFAP (in red) and nuclear (DAPI, in blue) staining of neonatal control (**E**) and mutant cortices without treatment (**F**) or mutant cortices after dorsomorphin (DM, **G**), S3I-201 (S3I, **H**), or dual (DM+S3I, **I**) treatment at E12.5.

**(J**) Quantitative analysis of GFAP-positive astrocyte numbers in the neonatal mutant cortex after treatment at E12.5.

Scale bar in (**A**), 100 μm for (**A** & **B**).

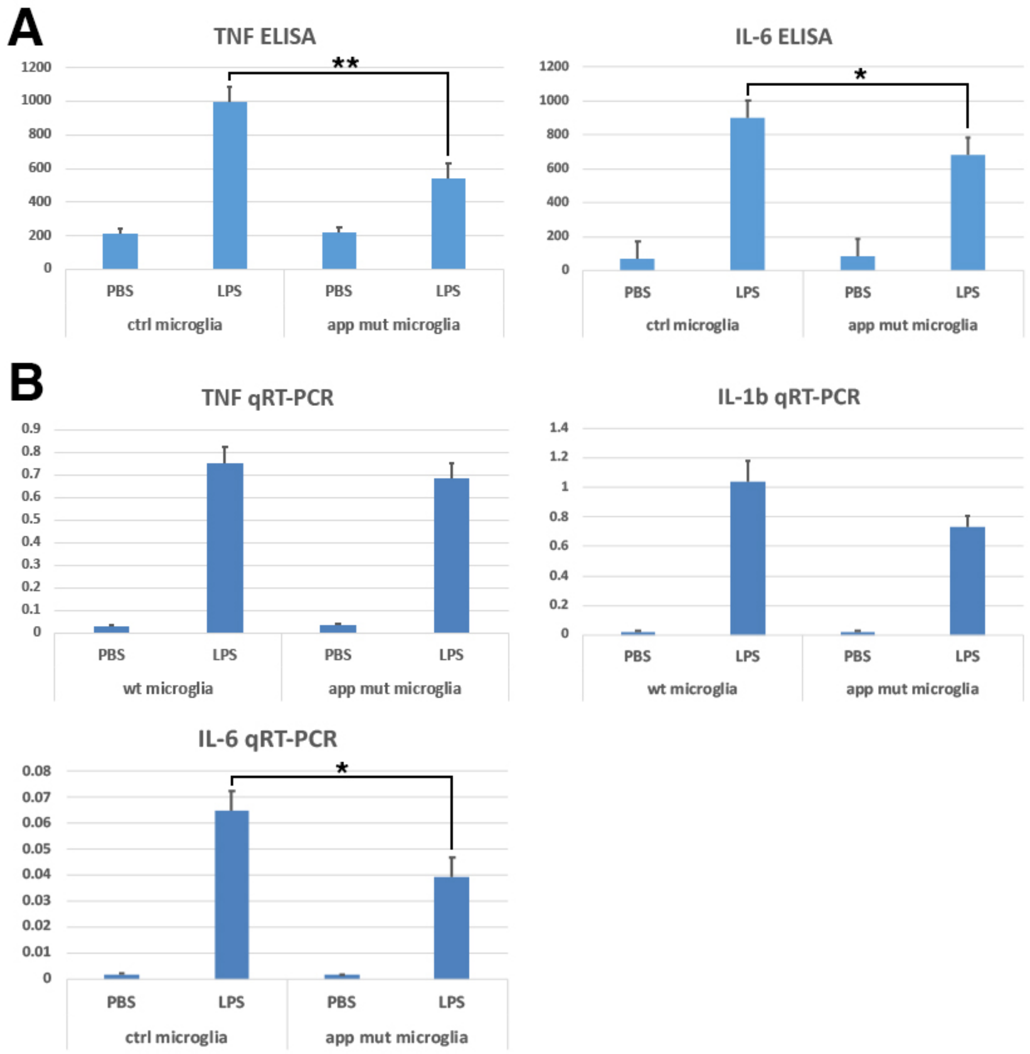


**Figure S6. Cytokine secretion and transcriptional induction in *app/cx3cr1-cre* mutant microglia.** Related to Figure 6.

**(A)** TNFα and IL-6 secretion (pg/ml) in control and *app/cx3cr1-cre* mutant microglia following overnight LPS stimulation. *, $P < 0.05$; **, $P < 0.01$; n = 9-13 each group.

(**B**) TNFα, IL-1β and IL-6 mRNA expression in control and *app/cx3cr1-cre* mutant microglia following overnight 3-hr LPS stimulation. *, $P < 0.05$; n = 6-7 each group.

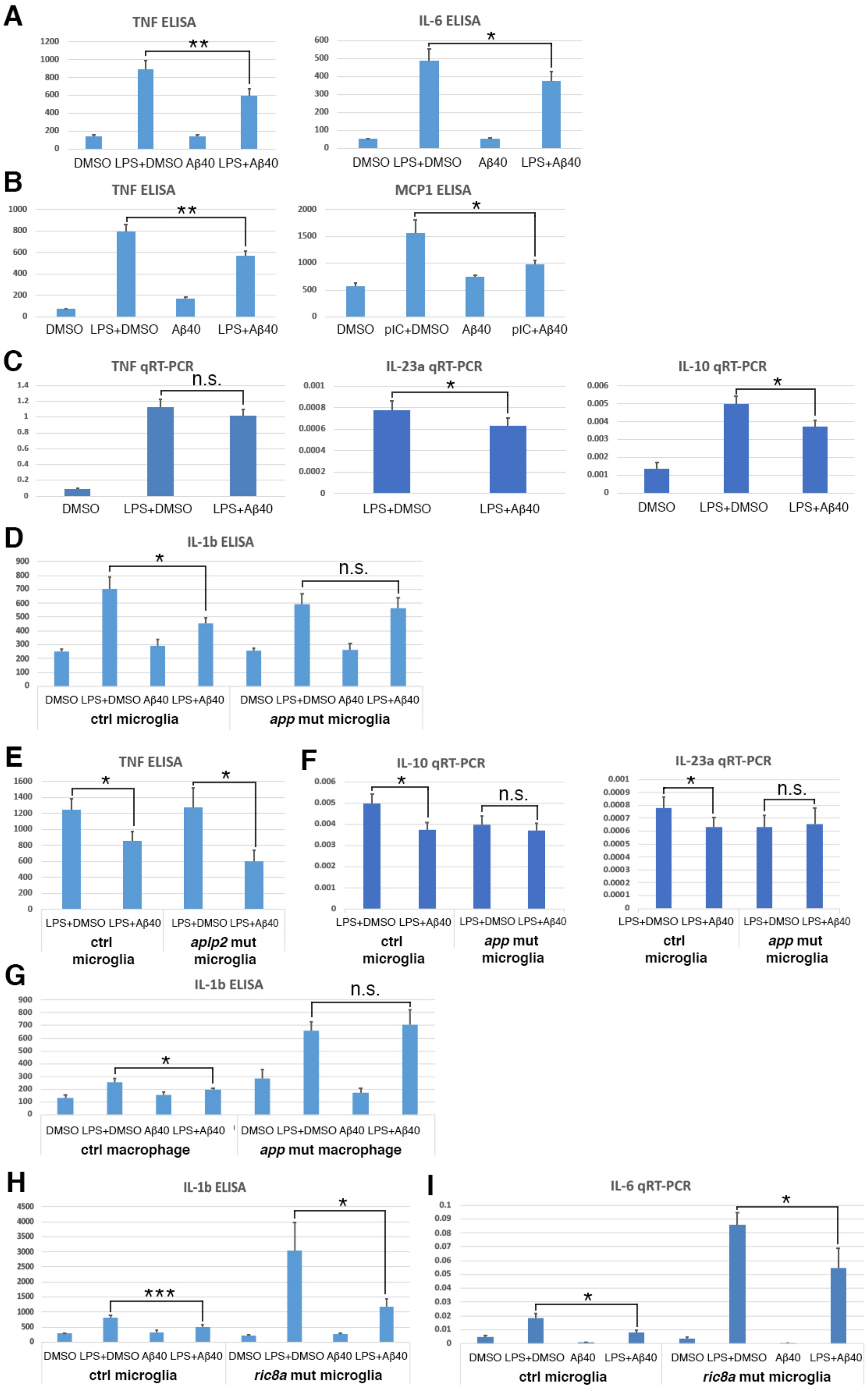
A
TNF ELISA
1200
1000
800
600
400
200
0
**
DMSO LPS+DMSO Aβ40 LPS+Aβ40
IL-6 ELISA
600
500
400
300
200
100
0
*
DMSO LPS+DMSO Aβ40 LPS+Aβ40
B
TNF ELISA
1000
800
600
400
200
0
**
DMSO LPS+DMSO Aβ40 LPS+Aβ40
MCP1 ELISA
2000
1500
1000
500
0
*
DMSO pIC+DMSO Aβ40 pIC+Aβ40
C
TNF qRT-PCR
1.4
1.2
1
0.8
0.6
0.4
0.2
0
n.s.
DMSO LPS+DMSO LPS+Aβ40
IL-23a qRT-PCR
0.001
0.0008
0.0006
0.0004
0.0002
0
*
LPS+DMSO LPS+Aβ40
IL-10 qRT-PCR
0.006
0.005
0.004
0.003
0.002
0.001
0
*
DMSO LPS+DMSO LPS+Aβ40
D
IL-1b ELISA
900
800
700
600
500
400
300
200
100
0
*
n.s.
DMSO LPS+DMSO Aβ40 LPS+Aβ40
ctrl microglia
DMSO LPS+DMSO Aβ40 LPS+Aβ40
app mut microglia
E
TNF ELISA
1600
1400
1200
1000
800
600
400
200
0
*
*
LPS+DMSO LPS+Aβ40
ctrl microglia
LPS+DMSO LPS+Aβ40
aplp2 mut microglia
F
IL-10 qRT-PCR
0.006
0.005
0.004
0.003
0.002
0.001
0
*
n.s.
LPS+DMSO LPS+Aβ40
ctrl microglia
LPS+DMSO LPS+Aβ40
app mut microglia
IL-23a qRT-PCR
0.001
0.0009
0.0008
0.0007
0.0006
0.0005
0.0004
0.0003
0.0002
0.0001
0
*
n.s.
LPS+DMSO LPS+Aβ40
ctrl microglia
LPS+DMSO LPS+Aβ40
app mut microglia
G
IL-1b ELISA
900
800
700
600
500
400
300
200
100
0
*
n.s.
DMSO LPS+DMSO Aβ40 LPS+Aβ40
ctrl macrophage
DMSO LPS+DMSO Aβ40 LPS+Aβ40
app mut macrophage
H
IL-1b ELISA
4500
4000
3500
3000
2500
2000
1500
1000
500
0
***
*
DMSO LPS+DMSO Aβ40 LPS+Aβ40
ctrl microglia
DMSO LPS+DMSO Aβ40 LPS+Aβ40
ric8a mut microglia
I
IL-6 qRT-PCR
0.1
0.09
0.08
0.07
0.06
0.05
0.04
0.03
0.02
0.01
0
*
*
DMSO LPS+DMSO Aβ40 LPS+Aβ40
ctrl microglia
DMSO LPS+DMSO Aβ40 LPS+Aβ40
ric8a mut microglia

**Figure S7. Aβ monomer effects on cytokine secretion and transcriptional induction in control and mutant microglial lineage cells.** Related to Figure7.

**(A)** TNFα and IL-6 secretion (pg/ml) in wildtype microglia following LPS stimulation in the absence or presence of Aβ40 (50nM). *, $P < 0.05$; **, $P < 0.01$; n = 25 each group for TNFα and 11 each group for IL-6.

(**B**) TNFα and MCP1 secretion (pg/ml) in wildtype microglia following LPS stimulation in the absence or presence of Aβ40 (500nM) from Genscript. Effects on IL-1β secretion in **Fig. 7B** was also performed with Genscript Aβ40. All other experiments in Fig. 7 were performed with ApexBio Aβ40. *, $P < 0.05$; **, $P < 0.01$; n = 5-7 each group.

(**C**) TNFα, IL-23, and IL-10 mRNA expression in wildtype microglia following LPS stimulation in the absence or presence of Aβ40 (400nM). *, $P < 0.05$; n = 6 each group

**(D)** IL-1β secretion (pg/ml) in control and *app/cx3cr1-cre* mutant microglia following LPS stimulation in the absence or presence of Aβ40. *, $P < 0.05$; n = 8-12 each group.

**(E)** TNFα (pg/ml) in control or *aplp2/cx3cr1-cre* mutant microglia following LPS stimulation in the absence or presence of Aβ40 (400nM). *, $P < 0.05$; n = 9-13 each group.

**(F)** IL-10 and IL-23 mRNA expression in control and *app/cx3cr1-cre* mutant microglia following LPS stimulation in the absence or presence of Aβ40 (400nM). *, $P < 0.05$; n = 6 each group.

**(G)** IL-1β secretion (pg/ml) in fresh unelicited control and *app/cx3cr1-cre* mutant peritoneal macrophages following LPS stimulation in the absence or presence of Aβ40 (400nM). *, $P < 0.05$; n = 12 each group.

(**H**) IL-1β secretion (pg/ml) in f control and *ric8a/cx3cr1-cre* mutant microglia following LPS stimulation in the absence or presence of Aβ40 (500nM). *, $P < 0.05$; ***, $P < 0.001$; n = 7-8 each group.

(**I**) IL-6 mRNA expression in control and *ric8a/cx3cr1-cre* mutant microglia following LPS stimulation in the absence or presence of Aβ40 (200nM). *, $P < 0.05$; n = 6 each group.
